# Population History and Admixture of the Fulani People from the Sahel

**DOI:** 10.1101/2024.06.22.600206

**Authors:** Cesar A. Fortes-Lima, Mame Yoro Diallo, Václav Janoušek, Viktor Černý, Carina M. Schlebusch

**Affiliations:** Human Evolution, Department of Organismal Biology, Evolutionary Biology Centre, Uppsala University, Uppsala, Sweden; McKusick-Nathans Institute and Department of Genetic Medicine, Johns Hopkins University School of Medicine, Baltimore, Maryland, USA; Archaeogenetics Laboratory, Institute of Archaeology of the Academy of Sciences of the Czech Republic, Prague, Letenská 1, 118 00 Prague, Czech Republic; Department of Anthropology and Human Genetics, Faculty of Science, Charles University in Prague, 128 01 Prague, Czech Republic; Palaeo-Research Institute, University of Johannesburg, Johannesburg, South Africa; SciLifeLab, Uppsala, Sweden; These authors contributed equally to this work

## Abstract

The Fulani people, one of the most important pastoralist groups in sub-Saharan Africa, are still largely underrepresented in population genomic research. They speak a Niger-Congo language called Fulfulde or Pulaar and live in scattered locations across the Sahel/Savannah Belt, from the Atlantic Ocean to Lake Chad. According to historical records, their ancestors spread from Futa Toro in the Middle Senegal Valley to Futa-Jallon in Guinea, and then eastward into the Sahel belt over the past 1500 years. However, the earlier history of this traditionally pastoral population has not been well studied. To uncover the genetic structure and ancestry of this widespread population, we gathered genome-wide genotype data from 460 individuals across 18 local Fulani populations, along with comparative data from both modern and ancient worldwide populations. This represents the most geographically wide-scaled genome-wide study of the Fulani to date. We revealed a genetic component closely associated with all local Fulani populations, suggesting a shared ancestral component possibly linked to the beginning of African pastoralism in the Green Sahara. Comparison to ancient DNA results also identified the presence of an ancient Iberomaurusian associated component across all Fulani groups, providing novel insights into their deep genetic history. Additionally, our genetic data indicate a later Fulani expansion from the western to the eastern Sahel, characterized by a clinal pattern and admixture with several other African populations north of the equator.

## Introduction

Fulani populations live in scattered areas across the Sahel/Savannah Belt (hereafter Sahel) with a population of around 25 million, exhibiting various traditions and lifestyles ^1^. They predominantly inhabit regions within West and Central Africa, including Adamawa, Kanem-Bornu, Futa-Masina, Futa-Jallon, and Futa-Toro, spanning eleven African countries: Mauritania, Senegal, Gambia, Guinea, Mali, Burkina Faso, Niger, Nigeria, Cameroon, Chad, and Sudan. Depending on the region, different terms are used for the Fulani people. The Hausa term “Fulani” is the most widely used, while the Wolof term “Peul” (or Pheul) was adopted by French and German speakers in the Middle Senegal Valley. Other terms include “Tukolor” (or Toucouleur, derived from Tekrur), “Toroobe” (from Islamic clerics), “Haalpulaar’en” (used by Pulaar-speakers), “Felaata” (used by Kanuri people), “Bororo” (to refer to Fulani cattle herders), and “Fulani Sire” (to refer to Town Fulani or the Hausa term “Fulani Gida,” which translates as “House Fulani”). More recently, the Fulfulde/Pulaar term Fulɓe (sg. Pullo) has been anglicized as Fulbe, which is increasingly used.

While the Fulani were traditionally considered nomadic pastoralists, raising mainly cattle, as well as goats and sheep, in the vast arid hinterlands of the Sahel/Savannah Belt, many have adopted a sedentary lifestyle ^2^. Groups of purely pastoral nomads are known by the Hausa name Mbororo’en (sg. Mbororo), but they call themselves Woɗaaɓe (sg. Boɗaaɗo). They keep zebu cattle, and between 45,000 to 100,000 individuals live in scattered camps in southern Niger, northern Nigeria, northern Cameroon, and adjacent areas of Chad and Burkina Faso ^3^. Today, a large portion of the Fulani comprises semi-nomadic or fully sedentary communities. These groups may have descended from former pastoralists, engaged in recent intermarriage with neighboring sub-Saharan African groups, or are the descendants of neighboring ethnic groups due to the so-called Fulanization process ^4^. Therefore, when collecting samples from Fulani communities, it is essential to consider their complex distribution and diversity to gain a comprehensive understanding of their population history in Africa.

The origin of the Fulani has been a long-standing debate ^5,6^. Their European morphological features combined with specific practices for appearance in females (e.g., tattoos, scarifications, decorations), as well as a moral code (called pulaaku) distinguishing them from neighboring communities, have given the impression that their ancestors came to West Africa from elsewhere ^7^. In addition, due to the strong cultural ties of the Fulani pastoralists to their cattle ^6,8,9^, which were not domesticated in Africa but in Southwest Asia ^10^, some scholars have suggested that the Fulani ancestors might have come from the Near East ^11^. However, other scholars have localized their putative homeland in the Nile Valley, considering ethnographic and historical records ^12,13^. Putative ancestors of the Fulani were also associated with Saharan rock art ^14,15^, interpreting some scenes in Tassili n’Ajjer (highlands in southern Algeria) as representations of Fulani rituals and ceremonies that survived millennia until recent times ^16^, but these conclusions have been later questioned ^17^.

Based on linguistic research, the Fulani language (called “Pulaar” or “Fulfulde”) belongs to the Atlantic branch of the Niger-Congo family, with the origin of this branch located in West Africa ^18,19^, where most Fulani populations live today. All language classifications attribute the Fulani language to the Niger-Congo family, deeply embedded in the western part of the Sahel belt ^20^. From a linguistic perspective, western Africa is the most likely origin of the languages spoken by the Fulani people. Currently, linguistic dialects in the Fulani language are divided into Pulaar in the west and Fulfulde in the east, which further includes approximately ten different subgroups (two in Pulaar and eight in Fulfulde) ^21^ that are closely related to languages from Senegal, such as Wolof and Serer ^22^.

Genetic studies of local Fulani populations or communities have become available more recently. One of the first studies ^23^ that focused on the fully nomadic groups of Fulani in Chad, Cameroon, and Burkina Faso showed that most of the Fulani mtDNA haplotypes (∼80%) were associated with West African ancestry, but a non-negligible amount (∼20%) was of West Eurasian or North African origin. These results were later confirmed in a large-scale study ^24^, which also included new mtDNA data from local Fulani populations in Mali and Niger. It was also shown that Fulani people have West Eurasian Y-chromosome haplogroups and that their mtDNA diversity is reduced compared to their Y-chromosome diversity ^25^.

If we consider lifestyle, all nomadic pastoralists in the Sahel are more likely than sedentary farmers to carry West Eurasian mtDNA haplogroups ^26^. Interestingly, nomadic pastoralists share mtDNA lineages belonging to haplogroups U5b1b and H1, which arose in sub-Saharan Africa after the dispersal of southwestern European populations around 8.6 thousand years ago (kya) ^27^. Further investigations have revealed the emergence of mtDNA sub-haplogroups specific to the Fulani, such as U5b1b1b and H1cb1 ^28^. Mitochondrial DNA studies of local Sahelian populations (both farmers and pastoralists) also revealed less gene flow between western Sahelian pastoralists (represented here mainly by Fulani) and their sedentary neighbors than the gene flow observed between eastern Sahelian pastoralists (represented here mainly by Arabic-speaking groups) and their sedentary neighbors ^29^. These observations are intriguing because Arabic-speaking populations arrived in the Sahel belt relatively recently ^30^, while the Fulani have been part of the western Sahel belt for a more extended period. However, in a study combining data from both uniparental genetic systems of numerous Sahelian populations, there was no significant population structure, as there was more genetic variation within Sahelian groups than between groups ^31^.

In contrast, autosomal diversity in the Fulani has been less investigated. Previously, data from microsatellite and insertion/deletion markers of Fulani participants from Cameroon revealed non-negligible non-African ancestry in the Fulani and genetic affinities with Chadic and Central Sudanic-speaking populations ^32^. Genome-wide studies have further confirmed Western Eurasian and North African genetic admixture in the Fulani of around 20% ^33–35^, which is consistent with the proportion identified in mtDNA studies ^23,29^. This non-sub-Saharan admixture in the Fulani gene pool has also been highlighted by analyses of the LCT gene, which has undergone positive selection in the Fulani ^34^. They have a high frequency of the –13,910T allele, also present among European populations as well as certain western Sahelian pastoralists such as the Moors and Tuareg ^36^. It has been revealed that not only this variant but also all surrounding haplotypes (∼2 Mb) are shared between the Fulani pastoralists from Ziniaré in Burkina Faso, European, and North African populations, suggesting admixture of the Fulani ancestors with a North African population. Due to the strong selective sweep, the level of non-sub-Saharan ancestry in the Fulani individuals carrying the – 13,910T allele in the vicinity of the LCT gene is at a high frequency compared to that of the alternative (ancestral) –13,910*C allele ^34^.

Whole genome data from a recent study ^75^ showed that the Eurasian (or non-sub-Saharan) component within the Fulani population might be much older and possibly related to the Green Sahara period (12,000–5,000 years BP), when the first cattle pastoralists appeared in North Africa. Subsequently, as a consequence of climate changes, these cattle herders, originally from the Green Sahara (possibly ancestors of contemporary Sahelian pastoralists, including the Fulani), moved westwards and southwards and admixed with other sub-Saharan African populations. This suggests that the Fulani genetic ancestry profile is very complex, mirroring the climate change in the Holocene.

To gain a better understanding of the genetic differentiation and population history of Fulani populations, we gathered a comprehensive dataset of 460 Fulani individuals (including 273 newly genotyped participants), representing a total of 18 local populations from 9 African countries across a geographic range stretching from the Atlantic Coast in the west to Lake Chad in the east (**Figure 1A**). To our knowledge, this survey constitutes the most comprehensive genome-wide genotype dataset of the Fulani distribution to date. Together with data from modern and ancient datasets of worldwide populations, we investigated the ancestral origins and genetic affinities of the Fulani. Our findings shed new light on specific migration, population structure, and the genetic differentiation between and within Fulani populations, potentially linked to their ancient pastoral history in the Green Sahara ^35–37^, which is in accordance with archaeological evidence dating to ∼8 kya ^38^.

**Figure 1.**
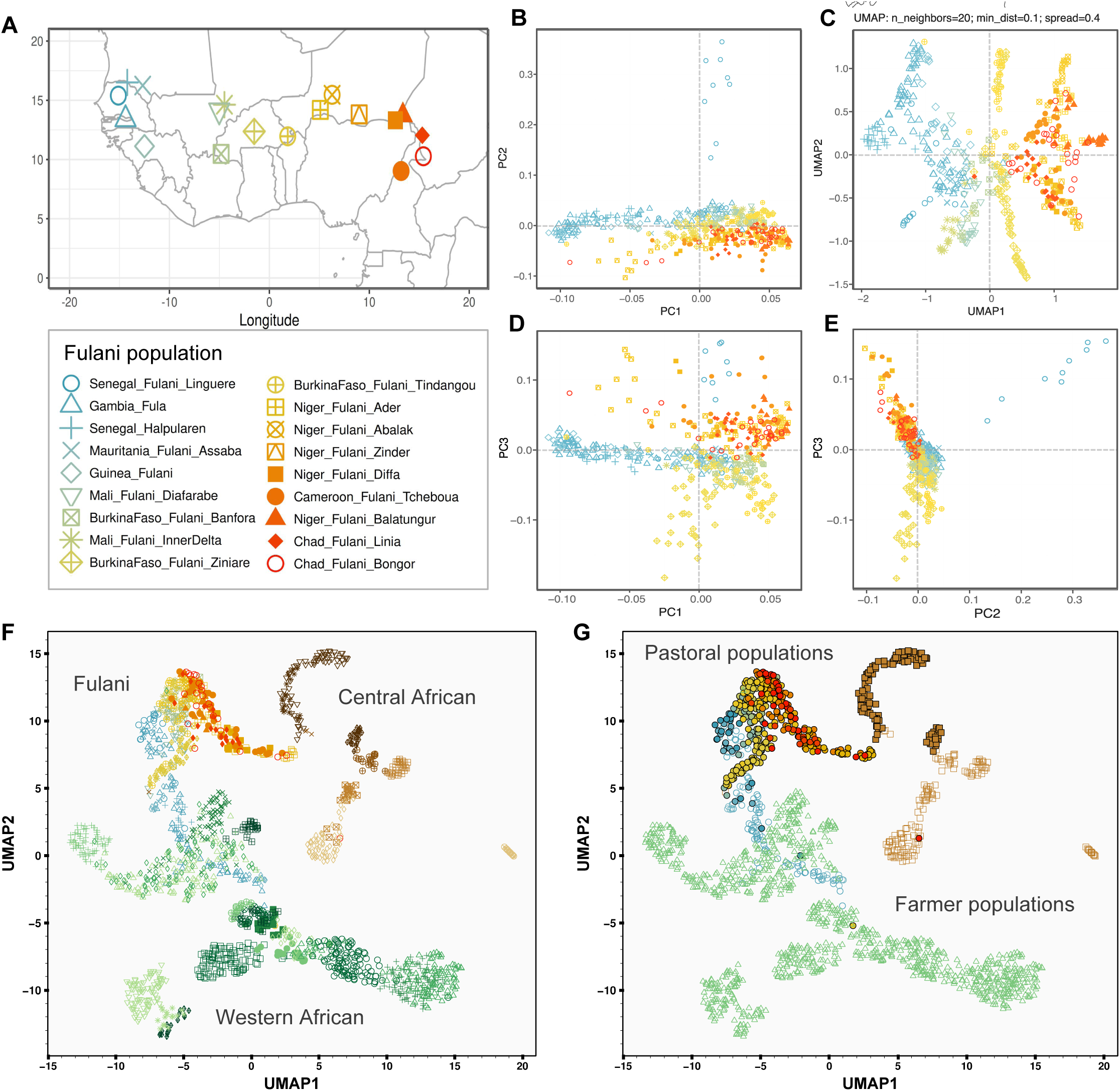
Dimension-reduction methods (DRM) used to explore the genetic diversity among studied Fulani populations. (A) Geographical distribution of all Fulani populations included in the Fulani-Only dataset (**Table S1**). (B) Principal component analysis (PCA) showing the distribution between PC1 and PC2. (C) PCA-UMAP approach combining the information of the first ten PCs. (D) PCA showing the distribution between PC1 and PC3. (E) PCA showing the distribution between PC2 and PC3. (F) PCA-UMAP combining the information of the first ten PCs estimated for Fulani and other local Western and Central African populations (parameters: n_neighbors=10 and min_dist=0.8). Colors and shapes of the markers of each population are the same that in **Figure S3A**. (G) PCA-UMAP plot highlighting with different color-coded shapes the subsistence context of Fulani (circles with the same colors than in F), western African non-Fulani (green triangles) and central African non-Fulani populations (brown squares). Subsistence strategies were indicated with empty shapes for farmer populations and filled shapes for pastoral populations (**Table S2**).

## Material and methods

### Sampling and genotyping

We collected samples from 419 Fulani volunteers (329 buccal swabs and 90 saliva samples) from 14 local Fulani populations. This collection was carried out during several years of fieldwork in Senegal, Mauritania, Guinea, Mali, Burkina Faso, Niger, Cameroon, and Chad (**Figure 1A** and **Table S1**). The study was approved by the Ethical Committee of the Charles University in Prague (approval number: 2019/12) and the Swedish National Ethical Review Authority (approval number: 2 2019-00479), and conducted according to the Declaration of Helsinki for medical research. At the SNP&SEQ Technology Platform (NGI/SciLifeLab Genomics, Sweden), DNA samples were genotyped on the Illumina Infinium H3Africa Consortium array (2,271,503 SNPs; using BeadChip type: H3Africa_2019_20037295_B1), designed to account for the large genetic diversity and small haplotype segments in African populations ^39^.

### Quality control and assembled datasets

We used PLINK v1.9 ^40^ to remove individuals with SNP-genotyping call rates equal to or lower than 85% and 116 individuals were removed (most were buccal swab samples). We used KING ^41^ to remove 30 individuals with a high probability of kinship up to the third degree. Our resulting dataset of 273 Fulani individuals was then merged with data of 187 Fulani individuals genotyped in previous studies using Illumina arrays: 74 Fula from Gambia ^37^; 54 Fulani from Burkina Faso ^34^; 23 Halpulaaren from Senegal ^35^; 25 Fulani from Guinea ^35^; and 13 Fulani samples from Burkina Faso, Chad and Niger ^33^ (**Table S1**). We performed quality control (QC) steps to keep only autosomal biallelic variants and individuals with high-genotyping rates (using PLINK as follows: --mind 0.15 -- geno 0.1 --hwe 0.0000001). After merging and QC, we obtained 1,141,817 SNPs and 460 individuals from 18 Fulani populations across 9 African countries in the “Fulani-Only” dataset (**Table S1**). We then merged the dataset with data from worldwide populations from previous studies, covering the genetic variation of reference populations in Africa, Europe and the Middle East ^33,37,42–46^. After merging and QC, we obtained 633,940 SNPs and 2691 individuals from 66 populations in the “Fulani-World” dataset (**Table S2**).

### Dimensionality reduction methods

To explore patterns of genetic affinities among all studied Fulani and comparative populations, we first used principal component analysis (PCA) ^47^ using smartPCA from the EIGENSOFT package ^47^. To avoid sample size bias due to our large sample of Fulani individuals, we employed the projection approach for PCA. First, we computed PCA for all the reference populations and a downsampled set of 36 randomly selected Fulani individuals from all studied Fulani populations, and then we projected onto the PCA the remaining 416 Fulani individuals. To combine the first 10 PCs, we used the PCA-UMAP approach ^48^. Results were visualized using the R package ggplot2 ^49^, and we also plotted PCs and geographical coordinates together. To analyse correlations between geography, subsistence and genetic variability as derived from PCA, we used linear models (ANCOVA) in R. To further test the effect of geography and subsistence on genetic distances among Fulani populations, we performed Mantel tests and multiple regression on distance matrices (MRM) using the ecodist R package ^50^. Among pairs of Fulani populations, we calculated the genetic distance matrix (*F_ST_*) using smartPCA, the geographical distance matrix using the geodist R package ^51^, and the subsistence matrix using codes of binary distances.

### Patterns of admixture and population structure

To investigate patterns of admixture and population structure, we performed clustering analysis using ADMIXTURE software v1.3.0 ^52^. For the Fulani-World dataset, we first used PLINK to remove SNPs under high linkage disequilibrium (LD) (as follows: --indep-pairwise 50 10 0.2). We obtained 2,691 individuals and 233,867 SNPs in the LD-pruned Fulani-World database. To avoid sample bias due to the large sample size of Fulani individuals, we employed the projection approach for ADMIXTURE analyses. We first computed PCA for reference populations and a downsampled set of 36 randomly-selected Fulani individuals from all studied Fulani populations and then we projected the remaining Fulani samples onto the PCA space. The same approach was applied to perform ADMIXTURE analyses in projection mode (-P) from K=2 to K=17. For each K, a cross-validation (CV) test was performed. The major mode for each K was visualized with bar plots using PONG ^53^, and pie chart plots were generated using custom R scripts. For visualization of the ADMIXTURE results using spatial interpolations, we applied the Kriging method and the grid-based mapping approach using Surfer software (Golden Software). To statistically test for admixture in Fulani populations, we used *f3*-statistics as part of ADMIXTOOLS ^54^. Worldwide reference populations included in the Fulani-World dataset were used as sources for admixture and Fulani populations as the target population. To investigate spatial patterns of migration and population structure across the Sahel belt, we used Fast Estimation of Effective Migration Surfaces (FEEMS) software ^55,56^.

### Comparison between modern and ancient DNA individuals

To investigate genetic links between Fulani populations and ancient DNA (aDNA) individuals, we merged the Fulani-World dataset with data from 91 aDNA individuals collected from previous studies (**Table S3**). We included three North African individuals ^57^ and 87 selected aDNA individuals included in the Allen Ancient DNA Resource (AADR) v54.1.p1) ^58^. After merging haplodized modern samples and pseudo-haplodized aDNA samples, we obtained 227,881 SNPs and 2,779 individuals in the “Fulani_aDNA-Modern” dataset. We used smartPCA to project ancient samples onto a background of present-day African populations (using “YES” for the following parameters: allsnps, lsqproject, newshrink, and killr2). ADMIXTURE analyzes from K=3 to K=8 were computed on the basis of the Fulani_aDNA-Modern dataset. The projection mode was used to project aDNA and modern Fulani individuals to a background of comparative modern populations that includes 36 selected Fulani individuals and worldwide populations following the approach explained above. To visualize ADMIXTURE results, we plotted the results for the K-group with the lowest cross-validation error using AncestryPainter v5.0 ^59^. We inferred the timing of admixture events using DATES v4010 ^60^. We inferred the time of the mixture by fitting an exponential distribution with an affine term using least squares.

### Admixture timing inference

To infer and estimate the dates of admixture events, we applied the admixture LD-based MALDER approach ^61^. For each Fulani population, we performed a multiple reference test using reference populations from various geographic locations, as well as the randomly selected sample of 60 Fulani individuals. The MinDis parameter, which represents the minimum genetic distance between a pair of SNPs to be considered, was set to 0.5cM ^61^. To convert the estimated duration of the generation into years from the dates deduced from the MALDER LD events, a calculation was applied: 1950 - (g x 29), where “g” represents the estimated number of generations and 29 the assumed length of one generation ^62^.

### Inferring demographic events among Fulani

To investigate demographic changes in Fulani populations, we estimated their effective population size (*N_e_*) in the last 50 generations using IBDNe analysis ^63^, based on the estimation of the rate of identity-by-descent (IBD) sharing between individuals of each population. We then converted estimated generations into years assuming a generation time of 29 years ^62^. To infer the age (*T_f_*) and the strength (*I_f_*) of putative founder events in studied populations, we applied ASCEND v8.6 ^64^ for each population included in the Fulani-World dataset using default settings. This approach measures the correlation in alleles sharing between pairs of individuals across the genome ^64^.

### Patterns of runs of homozygosity

To investigate population history and patterns of genomic inbreeding in the Fulani population, we calculated genome-wide runs of homozygosity (ROH) using a sliding-window approach implemented in PLINK following recommendations from ^65^. For each studied population, we used ROH segments shorter than 1.5Mb to calculate the sum of short ROH, and for segments longer than 1.5Mb we calculated the mean ROH size, sum of long ROH, total length of ROH and the genomic inbreeding coefficient (F_ROH_) using available R scripts (https://github.com/CeballosGene/ROH). We then investigate ROH segments of different lengths into six ROH length classes.

## Results

### Correlations between genetic, geographical and cultural diversity in the Fulani

Dimensionality reduction methods revealed a pattern of genetic diversity consistent with geographical differentiation, showing a west-east gradient among the studied Fulani populations (PC1 in **Figure 1B**) and PCA-UMAP (**Figure 1C**). On PC2, individuals from the Fulani population collected in Linguère (Senegal) separated out from other Fulani, while on PC3, Fulani populations in Burkina Faso differentiated from the Fulani in Niger, Chad and Cameroon (**Figure 1D–1E**). Heatmaps visualizing matrices of pairwise genetic and geography distances further highlighted the observed west-east gradient (**Figure S1**).

Subsistence of Fulani populations might also correlate with the observed genetic structure, where three western farmer Fulani (Fulani from Guinea, Fula from Gambia, and Halpulaaren from Senegal) have the lowest values on PC1 in contrast with Fulani that are pastoralists (**Figure S2A** and **Table S4**). To investigate this further, we tested the contribution of geography and subsistence as well as their mutual interactions. The model tested between longitude and subsistence explained 73% of the values estimated for PC1 (**Figure S2A**) and 72% for PC2 (**Figure S2B**) (both *P*-values were <0.001; **Table S5**). After removing all the outlier Fulani individuals from Linguere, PC2 evidenced a stronger significant correlation with longitude (F-statistic: 21.2, *P*-value= <0.001; **Figure S2C** and **Table S5**). In addition, Mantel tests between geography, subsistence and genetic distances were statistically significant (Spearman’s correlation= 0.35, *P*-value<0.001; **Table S6**); and when subsistence and geography were tested jointly using MRM, both factors were also highly significant (*P*-value= 0.007), with subsistence exhibiting the strongest effect (*P*-value= 0.001).

We then investigated patterns of genetic diversity in the Fulani together with worldwide populations using PCA (**Figure S4**) and by taking into account our large sample size of Fulani individuals using projected PCA (**Figure S5**). In both approaches, Fulani populations showed genetic affinities that match the observed west-east gradient (**Figures 2A–2B** and **S5**), with western Fulani closer to western African populations and other Fulani closer to central and eastern African populations. On the PCA-UMAP plots for only western and central African populations (**Figures 1F–1G** and **S6A**) and for all the studied populations (**Figures S6B–6C**), Fulani populations further showed the observed genetic patterns, which are also consistent with their type of subsistence due to the overlap between farmers from western Fulani and non-Fulani populations (particularly from Gambia), and the proximity between pastoralists from central Fulani and non-Fulani populations (**Figures 1F–1G**). Fulani populations were located together in a rotated V-shaped pattern (**Figures 2A** and **S4B**), which likely reflects putative admixture events involving different population sources (either between western and northern African sources or between central African and Eurasian sources).

**Figure 2.**
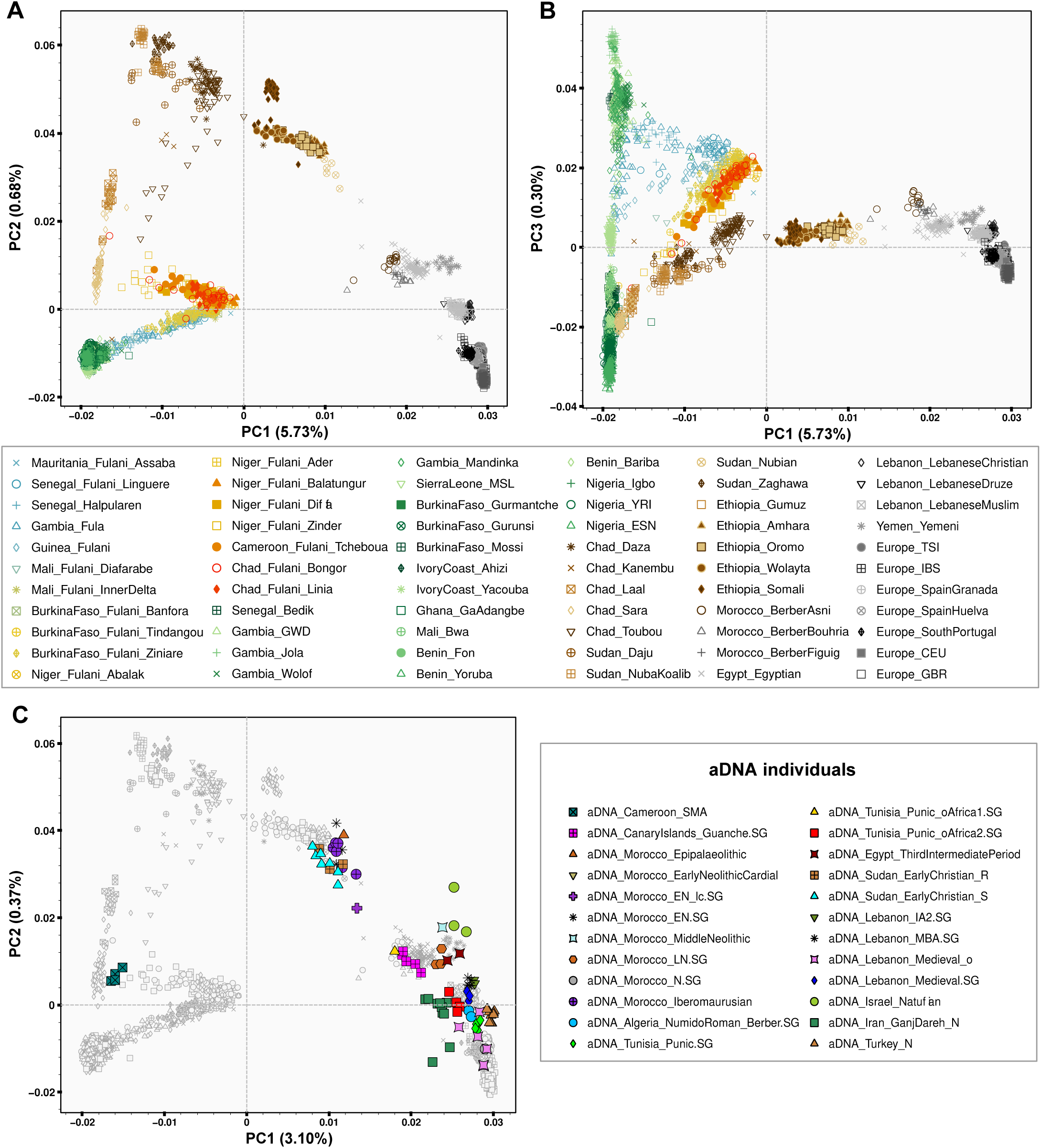
Genome-wide diversity of modern and ancient populations. (A) The first two principal components (PC1 and PC2) obtained using smartPCA for Fulani groups and all reference populations included in the Fulani-World dataset (**Table S2**). Downsampled Fulani set with the subsequent projection for the remaining Fulani samples was used to avoid sample size bias. (B) Figure showing PC1 to PC3 for all the populations included in the Fulani-World dataset (C) Figure showing PC1 to PC2 obtained using smartPCA to project 91 ancient samples (**Table S3**) onto the background of present-day African populations on the basis of the Fulani_aDNA-Modern dataset. Markers of ancient samples were filled with different colors, while markers of modern populations are in grey.

### Admixture and migration patterns in Fulani populations

Among Fulani populations, we observed different genetic contributions from a diverse range of ancestral sources in clustering analyses using the ADMIXTURE projection mode from K=2 to K=17 (**Figure S7**). At K=7, the K with the lowest value in the CV test (**Figure S8D**), we estimated a genetic ancestry predominant among all studied Fulani (green component; on average 45.6% SD=13.9%; **Figures 3A–3B, S7** and **Table S7**), and also among studied Moroccan Berbers at lower values (18.2% SD=1.74%; **Figure S8B**). Other components were also estimated among the studied Fulani, suggesting diverse degrees of admixture or ancestry sharing with Niger-Congo Atlantic (black component), Niger-Congo Volta (pink component), Nilo-Saharan Toubou (brown component), and Afro-Asiatic (dark purple component) populations (**Figure 3A–3B** and **S8**).

**Figure 3.**
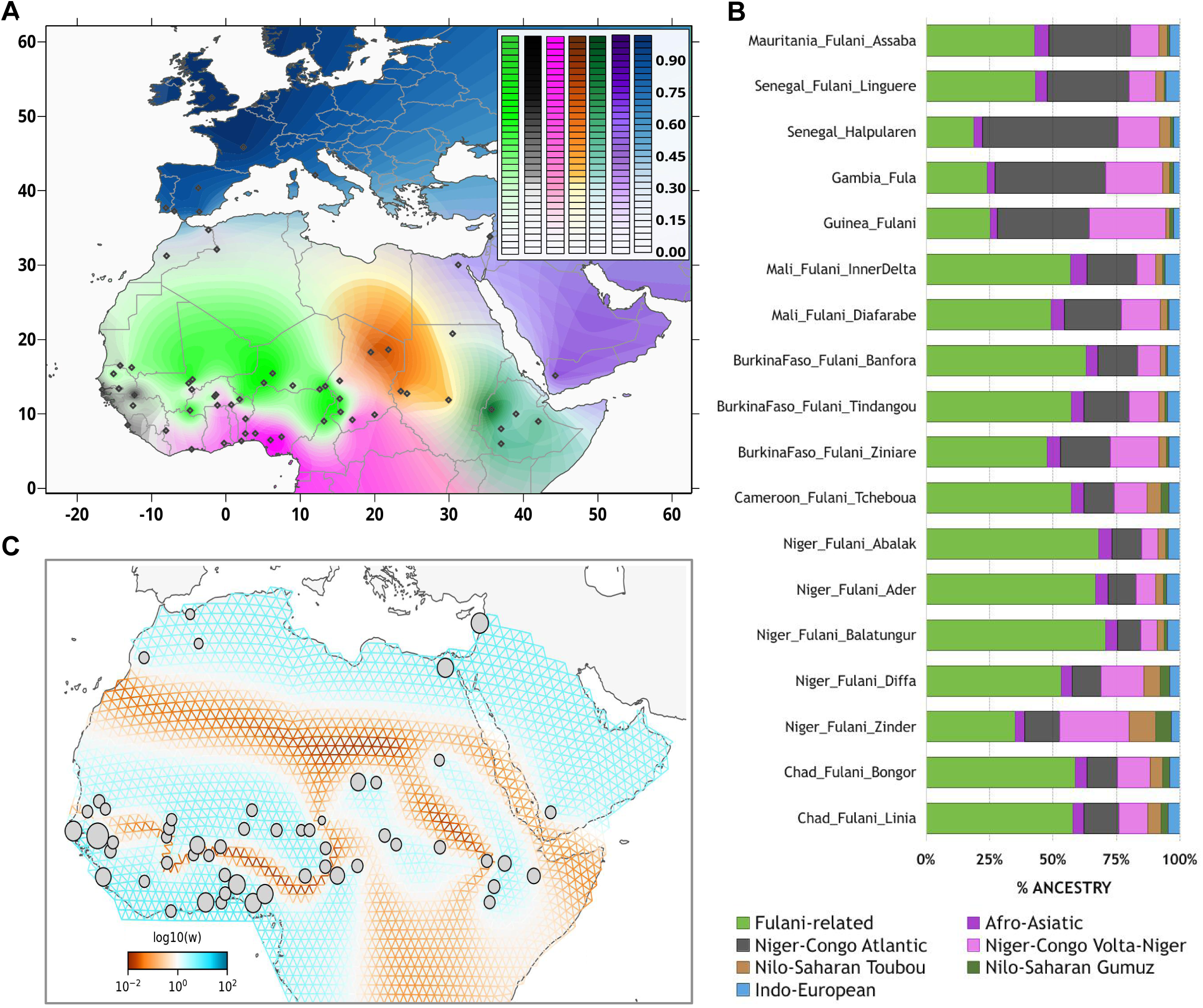
Genetic landscape of studied populations. (A) Pie chart plot based on ADMIXTURE results based on the Fulani-Wolrd dataset for K=7 raster plotted on a geographical map using the Kriging method. (B) Average values of the seven components estimated on ADMIXTURE results at K=7 for all studied Fulani populations. (C) Effective migration rates estimated using FEEMS. Figure showing fitted parameters in log-scale with lower effective migration shown using the orange pattern and higher effective migration shown using the blue pattern.

Consistent with previous studies ^34,35^, Fulani populations have a noticeable non-sub-Saharan African ancestry (blue component; range: 2.4–5.8% at K=7). Between the African sources, the strongest evidence of admixture inferred using *f3*-statistics was detected in western Fulani (from Mauritania, Gambia, Guinea, and Senegal; except for Fulani from Linguere) and central Fulani (from Zinder in Niger), involving one western African source and one northern African source (**Figure S9** and **Table S8**).

Estimated admixture events for each Fulani population successfully pinpointed 239 significant admixture-LD curves within 15 pairs of our weighted reference populations (**Figure S10** and **Table S9**). For the pairs of the selected Fulani–Senegal_Bedik and Fulani–Nigeria_Igbo, the chronology of the most recent episodes of admixture between Fulani and Niger-Congo-speaking populations showed events of admixture that took place between 7 and 25 generations ago (**Figure S10**). Interestingly, significant admixture-LD between Fulani and North African populations (e.g., selected Fulani and Moroccan Berber from Asni) highlights the depth and variability of these historical interactions (range: 49.5±5.9 to 74.5±5.7; **Table S9**). Other significant signals were observed involving Fulani and a Nilo-Saharan-speaking population from Central Africa (selected Fulani– Toubou range: 10.1±1.8 to 46.9±9.3), where Fulani from Linguere (Senegal) showed the oldest evidence of gene flow. We have also observed admixture events in the Fulani with a Nilo-Saharan-speaking population from East Africa (selected Fulani–Gumuz range: 40.9±7.1 to 8.0±1.0), where the Fulani from Assaba (Niger) showed the oldest evidence of admixture. Among all tested scenarios of admixture, western Fulani populations showed older evidence of admixture than eastern Fulani, which supports a west-east geographical cline with different episodes of admixture that occurred between the Fulani and local populations.

Besides the substantial patterns of admixture between Fulani populations and other groups, estimated effective migration surfaces showed population structure in Africa (**Figure 3C**), highlighting low migration rates between sub-Saharan and North African populations due to the presence of the Sahara Desert, in accordance with previous studies ^35,66^. Among sub-Saharan African populations, FEEMS analysis revealed a distinct genetic barrier along the western part, suggesting different geneflow patterns between Sahelian Fulani populations and western Niger-Congo populations. In addition, lower migration rates were estimated between central and eastern Sahelian groups, likely due to the presence of Lake Chad as another geographical barrier for gene flow. In contrast, patterns of high effective migration rates were observed among Nilo-Saharan speakers from Chad, Sudan, and Ethiopia.

### Comparisons between modern and ancient individuals

To further investigate the putative ancestors of the Fulani, we compared their genetic diversity with aDNA individuals (**Figure S3B** and **Table S3**). In the PCA (**Figure 2C** and **S11**), Fulani individuals are between modern and ancient individuals from sub-Saharan Africa and individuals from North Africa and Eurasia. Fulani from Cameroon and Niger have closer affinities with ancient Shum Laka individuals from Cameroon than western Fulani. Clustering analysis at K=6 evidenced a substantial presence of the Iberomaurusian (dark green) component among all Fulani groups (**Figure S12**). This component is also present in ancient Neolithic individuals from North Africa, in modern Berber groups from Morocco and certain population groups from Chad. The findings are in agreement with a recent study ^67^, while our study shows this component in a larger set of Fulani and other Sahelian and North African populations.

For ADMIXTURE results at K=8, Fulani groups receives their own (light green) component (**Figure S13**), that is also present among modern Moroccan Berbers (range: 5–9%). Interestingly, this Fulani-related component was also detected in ancient individuals from Algeria (9.6% in Berber-R10760.SG) ^68^ and Tunisian (10.3% in R11759.SG) ^69^, and five Guanche individuals from the Canary Islands (on average 8%) ^70^.

To infer admixture in the Fulani from putative ancestral sources, we selected Fulani populations from six different locations in the Sahel belt using two aDNA individuals from Cameroon as one source and aDNA samples from North Africa and the Canary Islands as the other source (**Table S10**). Estimated dates showed the oldest admixture event (510 ±208 generations; 14,800 ±6,026 years) in the Fulani from Abalak (Niger) between a Moroccan (5000 years; IAM) ^71^ and Western African (7000 years; SMA) ^72^ source. Also, older dates were inferred among the selected Fulani between those two sources (on average 254 generations) than with sources from Algeria, Tunisia, or the Canary Islands (**Table S10**).

### Demographic events and founder effects on Fulani populations

To shed additional light on the demographic histories of the Fulani, we applied three different methods. First, effective population sizes estimated using IBDNe showed demographic bottlenecks with minimum effective population sizes at around 25 generations ago (1225 CE; **Figures 4A** and **S14**), in agreement with a previous study using a limited number of Fulani individuals ^35^. The estimated *N_e_* showed more variation among western Fulani populations than among other studied Fulani (**Figure S14**). We also observed a population decline within the last 12 generations (circa 1600 CE) in the Fula (Gambia) and Fulani from Bongor (Chad), Tcheboua (Cameroon) and Banfora (Burkina Faso) (**Figures 4A**, **S14,** and **Table S11**).

**Figure 4.**
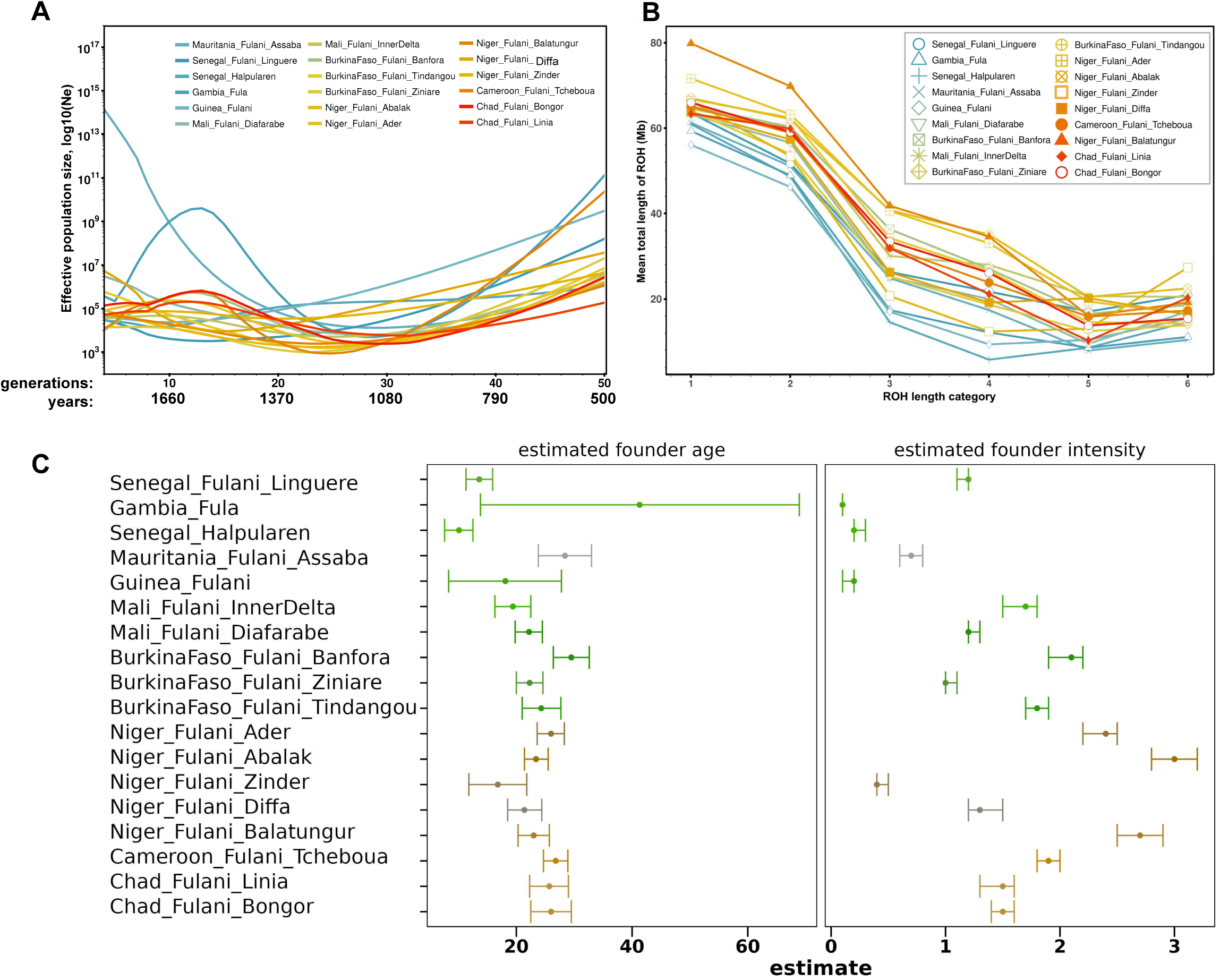
Demographic patterns among Fulani populations. (A) Effective population sizes (*N_e_*) among Fulani populations for the last 50 generations estimated using IBDNe. We converted inferred generations (g) to years using the following equation: 1950 – (g x29). (B) Categories of ROH length on the basis of the Fulani-Only dataset. Figure showing averages in each studied population and for each category of ROH length: class 1 for ROH length between [0.3-0.5Mb); clas 2 for [0.5-1Mb); class 3 for [1-2Mb); class 4 for [2-4Mb); class 5 for [4-8Mb); and class 6 for [8-16Mb). (C) ASCEND multiple reference test plots. Figure showing for each population the estimated founder age (*T_f_*, in generations before sampling) with standard error (SE) and estimated founder intensity (*I_f_*) with its SE.

Second, we shed new insight into the population dynamics and marriage customs of the Fulani using patterns of ROH. Among Fulani populations, the six categories of ROH lengths showed lower averages for western Fulani populations than for other studied Fulani (**Figures 4B, S15** and **Table S12**), with the highest values detected in Fulani from Niger (collected in Balatungur, Zinder and Abalak). In agreement with previous studies ^35,73^, Eurasian populations have the highest values for short categories of ROH and the total sum of short ROH, while western African populations have the lowest values (**Figures S15B, S16A** and **Table S12**). Therefore, higher values of the total sum of short ROH in Fulani populations than in Western African populations suggest gene flow with non-sub-Saharan African sources. Despite the similar values of Eurasian admixture in studied Fulani (**Figure 3B** and **Table S7**), we detected lower values of the total length of ROH among western Fulani (except for the Fulani in Linguere) than in other Fulani (**Figures S17B** and **Table S12**), suggesting different events of genetic isolation, inbreeding or demographic bottlenecks among the Fulani ^65^. In particular, the highest values of genomic inbreeding coefficient (on average F_ROH_= 0.052 ±0.037) and the total length of ROH segments (on average: 0.032 ±0.009) were estimated in the Fulani from Abalak in Niger (**Figures S17–S18**).

Third, we investigated the timing and intensity of founding events in the Fulani. In contrast with comparative populations, ASCEND results showed relatively recent founder events (on average, *T_f_*= 23 generations) of different intensities (range: 0.1–3) among Fulani populations (**Figures 4C S19–S20**, and **Table S13**). Significant founder events were observed in five Fulani populations (two from Mali (Inner Delta and Diafarabe); two from Niger (Ader and Abalak); and one from Banfora in Burkina Faso), suggesting a strong correlation in allele sharing between pairs of individuals in each population where the estimated ages (range: 16.8–29.5 generations), intensities (range: 1.8–3.0%) and NRMSD values (range: 0.028–0.039) are in agreement with all the required thresholds defined by Tournebize et al ^64^. The highest estimated founder intensity (*I_f_*= 3.0%) was inferred in the Fulani from Abalak, suggesting a more significant reduction in genetic diversity and an increased probability of a small founding population of this population.

## Discussion

### Genetic and cultural diversity of the Fulani

The Fulani people are one of the largest pastoral groups in Africa, known for maintaining diverse subsistence strategies ranging from fully pastoral to agro-pastoral and, in some cases, agricultural. This large nomadic group is distributed in scattered locations across sub-Saharan Africa, and their widespread presence and concentration in the Sahel/Savannah belt likely mirror the geographic origins of their ancestors ^2^. The originally pastoral nomadic lifestyle of the Fulani, along with their physical features ^74^, has often led neighboring communities to perceive them as transient. This perception has fueled the misconception that the Fulani are in perpetual migration and originated from elsewhere.

This study gathered genomic data from a comprehensive sample of 18 local Fulani populations, representing their complex gene pool across the region. Dimensionality reduction methods applied to the studied Fulani populations, both alone and in comparison with worldwide populations, highlighted genetic diversity that can be explained by a clinal pattern from the western through central to the eastern regions of the Sahel belt (**Figures 1–2**). This west-east differentiation aligns with the division of the Fulani linguistic dialects (Pulaar and Fulfulde) and shows significant correlations between genetic and geographical distances, as well as with the subsistence strategies of the Fulani (**Tables S4–S6**). This suggests that cultural factors might have contributed to their current genetic landscape. Genetic contributions to the Fulani were also inferred from populations belonging to the Atlantic and Volta-Niger branches of the Niger-Congo linguistic family. Linguistically, Fulfulde is part of the Atlantic branch of the Niger-Congo family ^20^, sharing affinities with Wolof and Serer. In accordance with linguistic records ^20^, clustering and f3-statistics analyses showed that western Fulani have more genetic affinities with western sub-Saharan African populations, while more Nilo-Saharan-related ancestry was detected in Fulani populations from Cameroon, Niger, and Chad (**Figures 3A–3B** and **Tables S7–S8**).

### Population history during the Green Sahara

Our large sample size of Fulani individuals representative of their wide distribution in Africa allowed us to gain new insights into their population structure. By using projection approaches, we addressed potential biases due to large sample sizes. Dimensionality reduction methods and clustering analyses using the projection mode depicted the complex patterns of admixture in the Sahel belt, which are in agreement with previous studies ^35,75–77^. Clustering analysis also identified a genetic component that is predominant in all studied Fulani populations (**Figure 3B**), reflecting their shared ancestry. This genetic component has a large distribution across western and central regions of the Sahel belt (from Senegal to Chad; **Figure 3A**), and in lower frequencies in northwestern Africa (Moroccan Berbers). This distribution is also consistent with the observed effective migration rates, where low effective migration rates were observed around the area of the estimated Fulani-related component (**Figure 3C**).

The comparisons of present-day Fulani and aDNA individuals from the Near East, North Africa, and sub-Saharan Africa revealed that the ancient Iberomaurusian component is present in all current-day Fulani groups, as well as in Berber populations from North Africa and certain populations from Chad (**Figure S12**). The clustering results in **Figure S12** suggest that the ancestral sources of the Fulani might have been a North African population (related to ancient North African Neolithic groups and current-day Berbers) and a West African population (related to current-day Gambian or Senegalese populations). Our admixture dates, using Early Neolithic individuals (from IAM, Morocco) and Early Stone to Metal Age individuals (from Shum Laka, Cameroon) as sources, indicated the oldest admixture dates in the Fulani (from Niger) around 14.8 kya (**Table S10**), possibly reflecting ancient contact between sub-Saharan and North African groups. Average dates of 254 generations (7.4 kya) were inferred among the Fulani with these two sources as parental groups, which falls within the Green Sahara period. The Green Sahara period was characterized by significantly higher rainfall than before, transforming deserted areas into fertile lands, and enabling rapid human population growth. This likely facilitated contact between the Fulani’s North African ancestral source (possibly already practicing nomadic pastoralism) and sub-Saharan populations ^78,79^.

The Sahara is the largest open-air museum of rock art, created initially by hunter-gatherers and later by pastoralists, featuring the so-called bovidian paintings that clearly show the introduction of Near Eastern domestic animals to Africa. The first presence of cattle is reported in Ti-n-Torha in the Acacus (7430 ± 220 years ago) ^80^, and along with bovine remains in northern Chad ^81^, this suggests the presence of pastoralists at least ∼7 kya ^82,83^. Both cattle and small livestock, such as goats and sheep, were introduced to the Green Sahara and gradually adopted by local hunter-gatherer groups ^81,84^. Around 2 kya, South Asian humped zebu was introduced via Arabia into Africa, and present-day Fulani incorporated this breed into their pastoral economy ^10^. Ancient contacts between populations in the Lake Chad Basin and Berbers were also suggested by the study of the L3e5 mtDNA haplogroup ^85^, and by the study of whole genomes of present-day Fulani ^67^.

Future studies generating aDNA data from individuals in the Sahara and Sahel belt will provide further insights into the historical distribution of the Fulani ancestors, their past migrations, and interactions with North African groups.

### Recent admixture events in the Fulani

Genetic contributions from Nilo-Saharan speakers (Toubou and Gumuz) in Fulani from Cameroon, Niger, and Chad suggest that the Fulani ancestors could have received gene flow somewhere in the central and eastern regions of the Green Sahara, where they interacted with ancient Nilo-Saharan peoples, possibly the Aquatic Civilization ^86^. These contacts could have been unidirectional since there are minimal contributions of the Niger-Congo component in modern Nilo-Saharan speakers in both the central and eastern Sahelian regions. It has been proposed ^87^ that the homeland of Niger-Congo and Nilo-Saharan languages was between the Maghreb and the Nile Valley during the final phase of the Late Pleistocene (20–12 kya), and their language diversifications accelerated as populations expanded within the Green Sahara at the beginning of the Holocene (12–10 kya). These observations highlight the important role of ancestral sub-Saharan sources in shaping the Fulani genetic heritage.

The estimated non-African genetic component in the Fulani gene pool coming from Near Eastern (on average 18%) and European (7%) sources is consistent with previous studies ^33–35^. Likely, the period this gene flow is connected to back-to-Africa migrations after the last glacial maximum (LGM) ^88^. Indeed, modern North Africans have genetic affinities to both West Eurasians (Europeans and Near Easterners) that reveal clinal patterns due to the continuous back-to-Africa migration(s) ^89^. Interestingly, Near Eastern Late Paleolithic and Neolithic populations also show a high level (∼44%) of Basal Eurasian ancestry ^90^, which has been formed in the Late Pleistocene refugium of the Arabo-Persian Gulf without admixture with Neanderthals ^91^. It is possible that one or more populations from the Near East migrated to the Maghreb already in the pre-agricultural period as hunter-gatherers ^92,93^, as already shown by analyses of aDNA extracted from Iberomaurusian skeletons ^94–96^.

By examining the decay of admixture LD in the Fulani populations with multiple population sources through MALDER, we have further highlighted the timing of historical admixture events between the Fulani ancestors and various genetic contributors. The observation of the North African admixture interval in the Fulani between 75 and 50 generations ago suggests that contact occurred throughout the first millennium AD, consistent with previous observations ^34^. That might be the last major admixture event in the Sahara as the southward progression of the desert and drying of the Saharan lakes around ∼3 kya ^97^ displaced various populations to more southerly regions with more water sources and more favourable conditions.

### Patterns of genetic isolation and population expansion in the Fulani

Our results using clustering and ROH analyses show higher genetic diversity among the studied Fulani than in their neighbouring Sahelian populations, except for western Fulani populations with low patterns of ROHs. Among studied populations, the highest values for long ROH categories were observed within Fulani populations, in particular for categories 4 and 5 (**Figure S15B** and **Table S12**), suggesting genetic drift in these populations. Among Fulani populations, regional differences were detected from the Abalak in Niger to Halpularen in Senegal (**Figures S15–S18**), suggesting different demographic events between Fulani populations across the west-east pattern. The highest values of genomic inbreeding coefficient in the Fulani from Abalak (Niger) suggest higher genetic isolation and likely a demographic bottleneck than in other Fulani populations (**Figures S18** and **Table S12**). Estimated effective population sizes showed different events of population decline and expansion among Fulani, highlighting different demographic events since the founding of those populations in different regions, in agreement with previous studies ^35,67^.

Besides the close geographical distribution of the Fulani and sub-Saharan populations, the highest frequencies of the Fulani-related genetic component are observed among Berbers (**Figure S8**). This suggests a shared population history between these two groups, possibly since their cohabitation during the last African Humid Period (AHP) ^98^, but might also be due to historically documented contacts with Berbers of Znaga in recent times. We can also notice Berber linguistic influences as indicated by correspondences and the influence of the Fulani language on local names in the Mauritanian regions of Brakna and Tagant ^99^. Nonetheless, the higher occurrences of this component among the Fulani of Niger or Chad suggest a more complex scenario regarding its origin in the eastern part of the Sahara, from where pastoralism has spread to the south and west.

We identified founder events in Fulani populations around 25 generations ago (**Figure 4C**). This is also supported by the IBDNe results (**Figure 4A**), which indicate expansion that occurred within the Fulani populations shortly after that time and these results are generally consistent with the observations from previous studies ^23,31,35^. This can be explained by environmental factors resulting from the wet phase that took place in the Sahel from 700 to 1400 AD. In Senegal River Valley circa 1300–1350 AD, the regional economy became progressively pastoral ^97^, which could have contributed to the spread of the Fulani populations to Futa-Jallon and further east via the Inner Delta of Niger to Lake Chad Basin as documented historically ^100^. The flourishing trans-Saharan trade during the last 500 years would have played a pivotal role in facilitating the development of extensive trade networks for the Fulani and the accumulation of wealth among their communities. Trans-Saharan caravans were often a place for the exchange of ideas, cultures and knowledge, which allowed some Fulani pastoralists to further participate in cultural and biological exchanges ^34^. Nevertheless, the majority of the Fulani population remained faithful to pastoralism (an archaeologically almost invisible lifestyle) ^2^.

## Conclusions

In summary, the observed genetic differences between local Fulani populations following a west-east pattern reflect their unique genetic history, shaped by interactions with different local groups and various demographic events. Our analyses revealed that subsistence strategies, along with geographical patterns, significantly influenced the observed diversity among local Fulani populations. Comparisons between modern and ancient DNA data allowed us to infer novel evidence of population structure and gene flow over time, identifying the ancient Iberomaurusian component in all Fulani groups. These findings indicate that the Fulani genetic ancestry is complex, with contributions from both North African and West African sources, and highlight the impact of historical migrations and climate changes in shaping their genetic landscape. This study addressed long-standing questions about the ancestral origins of the Fulani and provided new insights into their population structure, migration patterns, and admixture within the African continent.

## Acknowledgements

We thank the volunteers who participated in this study. We thank Mario Vicente and Francisco Ceballos for helpful discussions. The genome-wide SNP data computations were made possible by the National Academic Infrastructure for Supercomputing in Sweden (NAISS) resources provided at UPPMAX (project numbers: naiss2023-22-463 and naiss2023-22-464), which were partially funded by the Swedish Research Council through grant agreement no. 2018-05973. Additionally, computational resources were provided by the e-INFRA CZ project (ID:90254), supported by the Ministry of Education, Youth and Sports of the Czech Republic.

## Data availability

Novel SNP array genotype data of Fulani populations generated in this study will be made available through the European Genome-phenome Archive (EGA) data repository (EGA accessory numbers: TBD; data will be available online prior publication). C.M.S. was granted data access to genome-wide genotype data deposed by the APCDR AGV Project (EGA accessory number: EGAS00001000959) and the EUROTAST Project (EGA accessory number: EGAS00001002535).

## Code availability

Data analysis scripts and R scripts for plotting used in this project are available online on GitHub (https://github.com/vjanousk/h3a-fulani), as well as interactive plots and Python scripts for plotting (https://github.com/Schlebusch-lab/Sahel_study).

