## Supplementary material for "Population History and Admixture of the Fulani People from the Sahel": Figure S

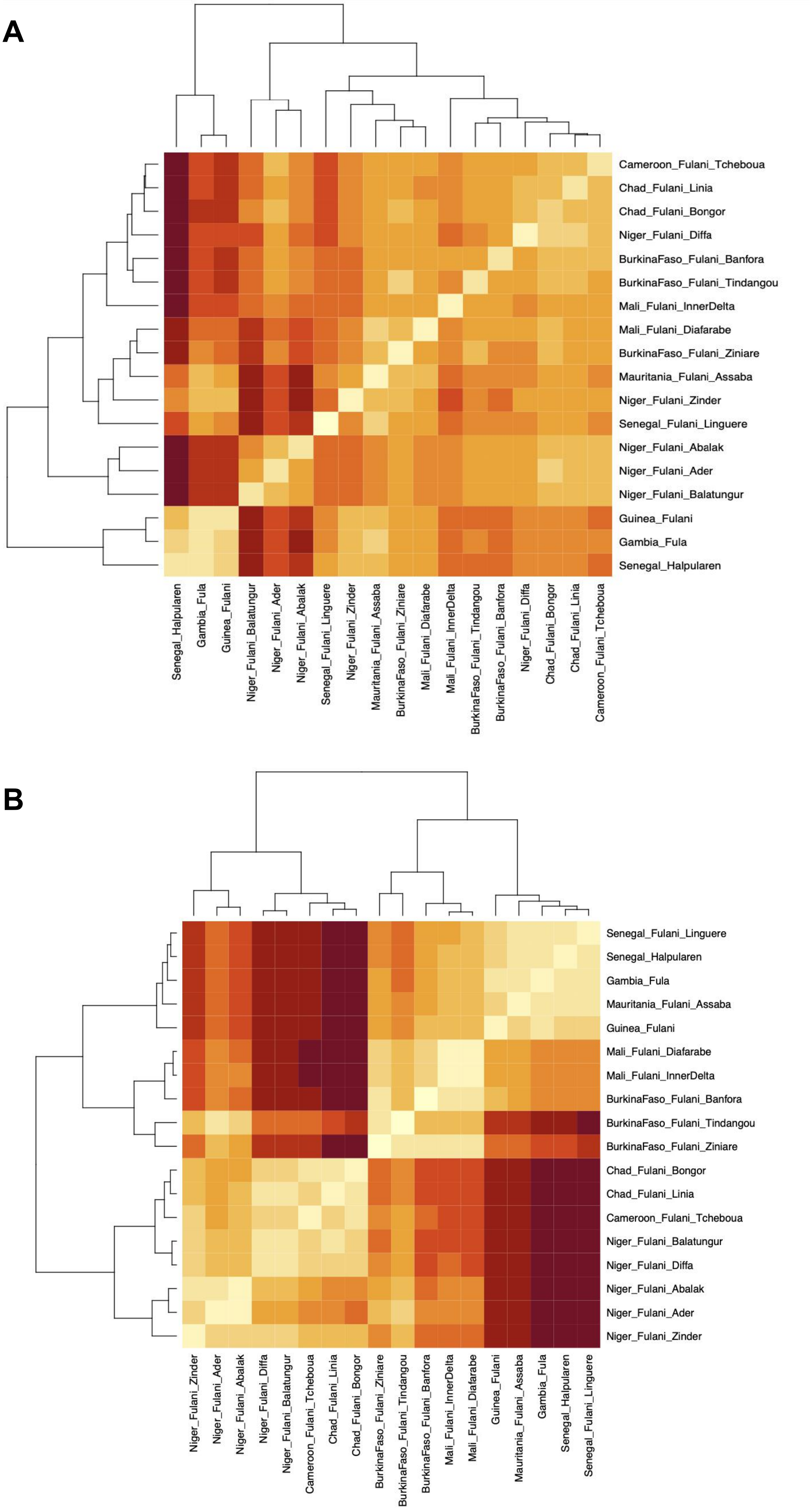


**Figure S1. Distance matrices used in Mantel tests.** (A) Pairwise genetic distances ($F_{ST}$) between the populations included in the Fulani-Only dataset and calculated using smartPCA. (B) Pairwise geographical distances based on the approximate sampling location of each studied population.

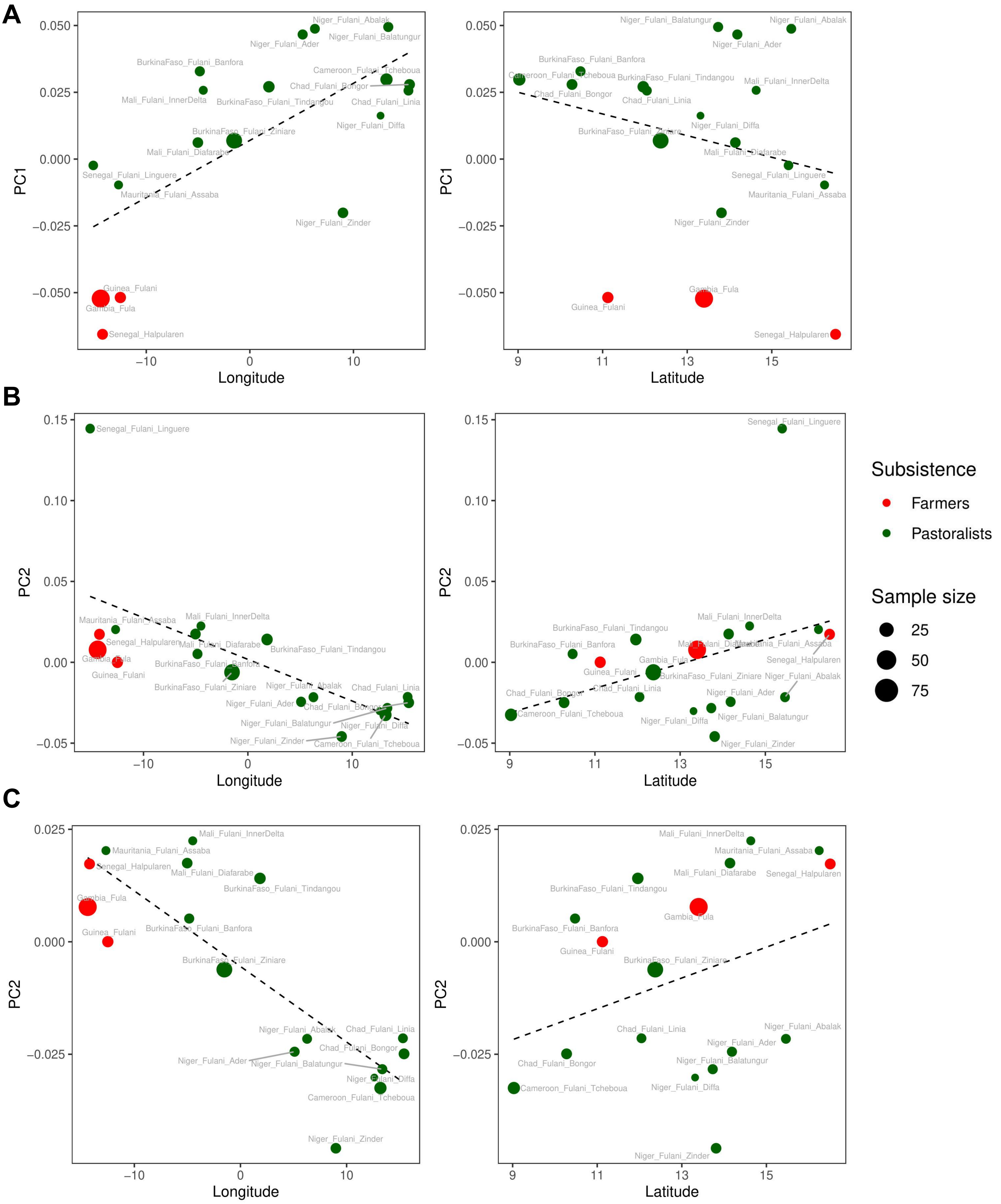


**Figure S2. Correlation of principal components (PC) with geography and subsistence.** (A) Correlation plots between PC1 and longitude (left) and latitude (right). (B) Correlation plots between PC2 and the longitude and latitude including all studied Fulani individuals; and (C) after removing the Fulani_Linguere population from Senegal. Color-codes correspond to the subsistence, and the size of the dots correspont with each population size. Estimated values were included in **Table S4** and the results of the tested models in **Table S5**.

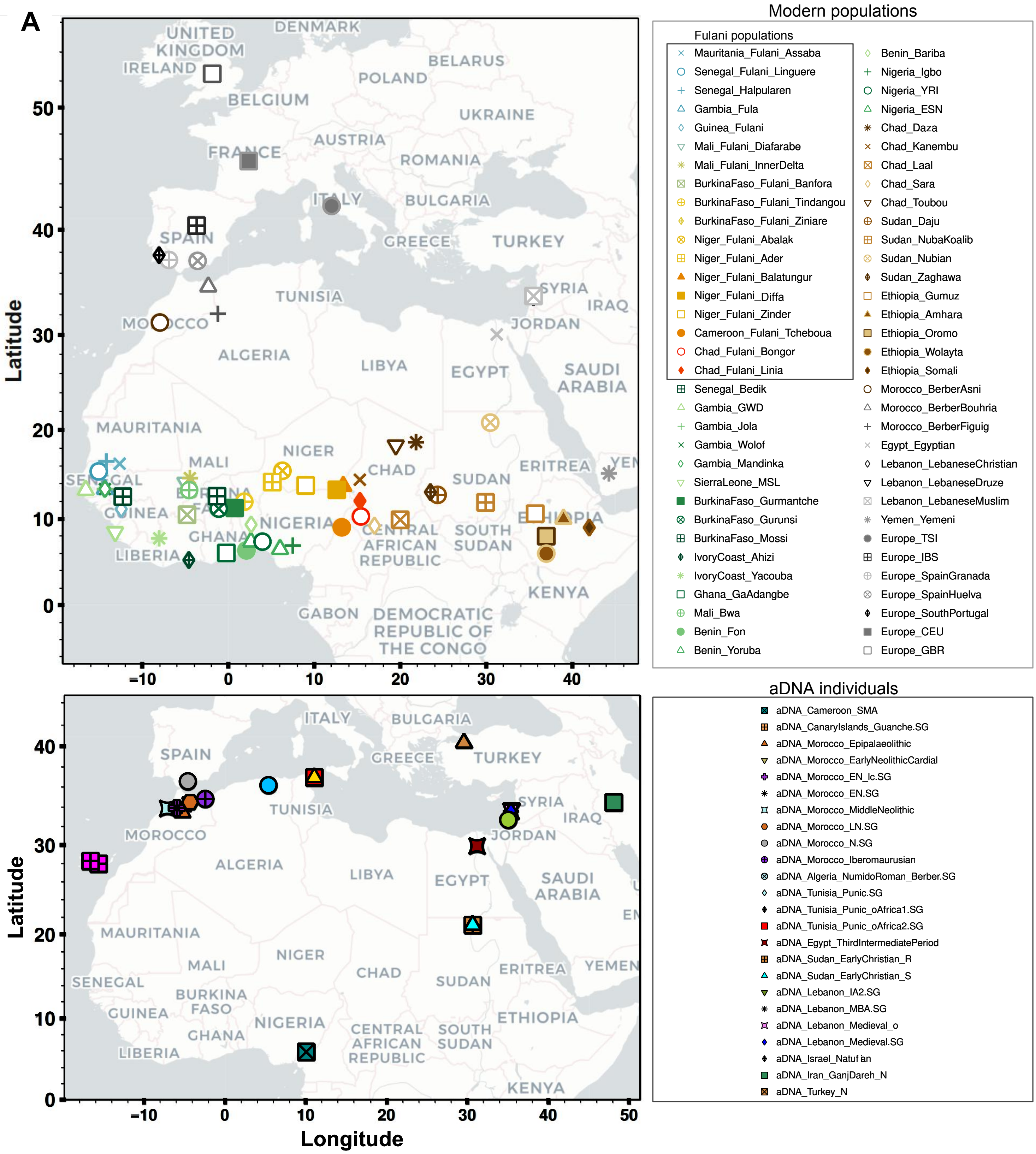


**Figure S3. Distribution of modern and ancient populations included in this study.** (A) Figure showing all the populations included in the Fulani-World dataset. Details of each studied population were also included in **Table S2**. (B) Figure showing all the populations included in the Fulani_aDNA-Modern dataset. Details of each studied population were also included in **Table S3**. To better visualize the results of each studied population, we included interactive plots in Github (https://github.com/Schlebusch-lab/Sahel_study).

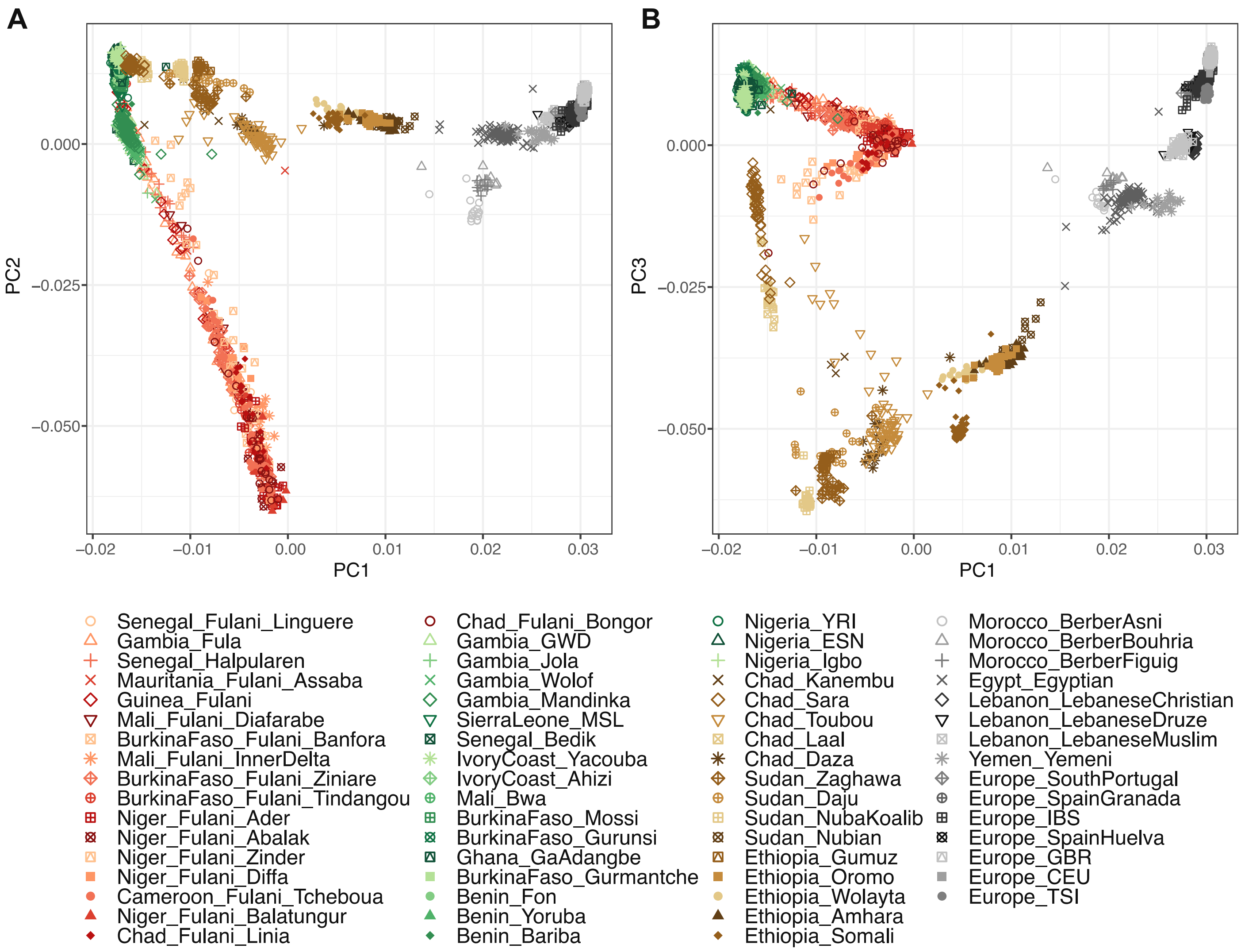


**Figure S4. Un-projected PCA for Fulani and comparative populations.** (A) Figure showing un-projected PCA between PC1 and PC2; and (B) between PC1 and PC3. PCA was performed for all the populations included in the Fulani-World dataset, and without downsampling Fulani individuals and subsequent PCA projection for the remaining Fulani individuals like in **Figure S5**. Details of each studied population were also included in **Table S2**.

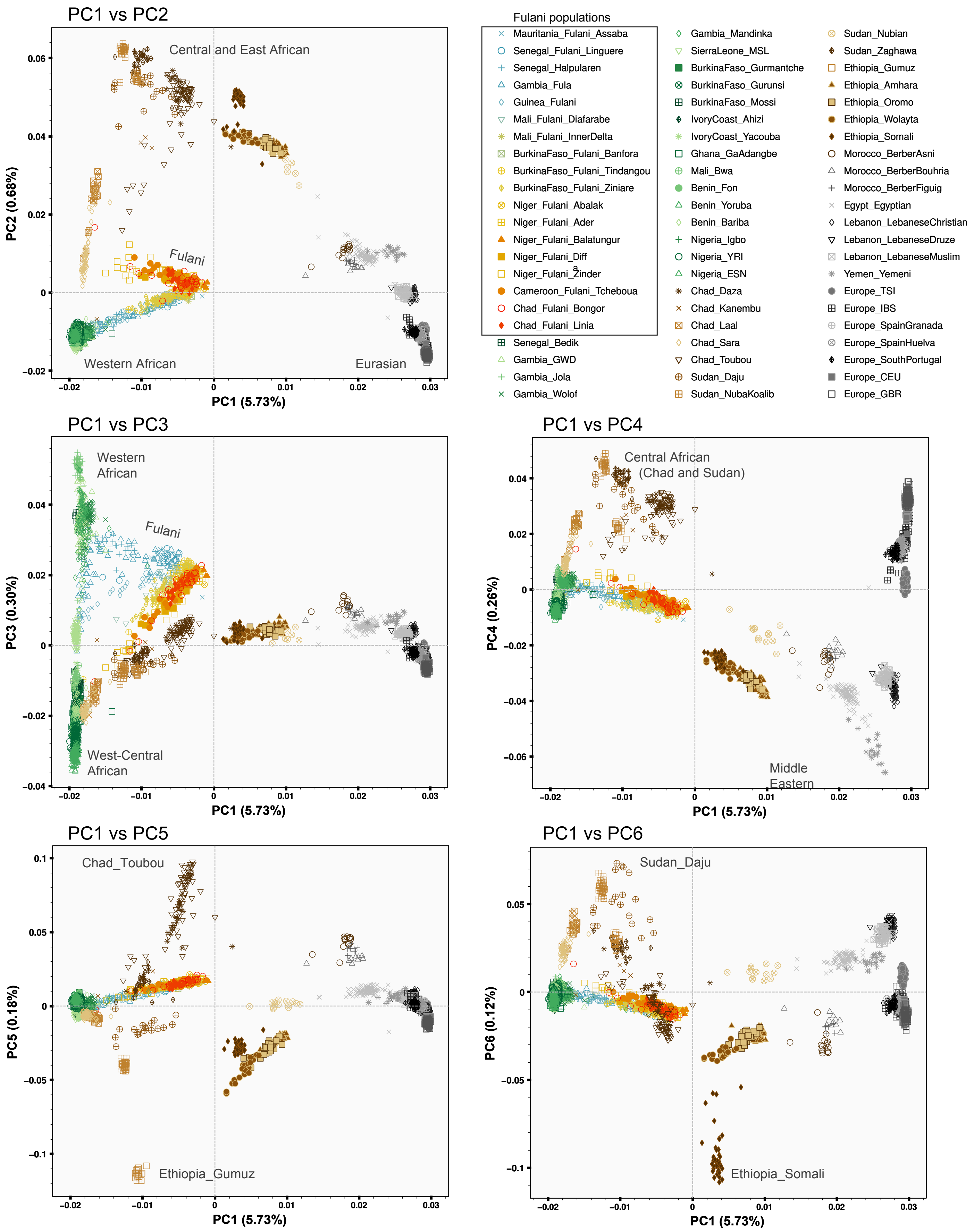


**Figure S5. Genome-wide diversity of the Fulani and worldwide comparative populations**. Figure showing PCA for the first six principal components (PC) estimated for all the populations included in the Fulani-World dataset. To avoid sample size bias (**Figure S4B**), we first computed PCA for reference populations and a downsampled set of 36 randomly-selected Fulani individuals from all studied Fulani populations, and we then projected onto the PCA the remaining Fulani samples. Geographical locations of the studied populations were included in **Figure S3A**. To better visualize the results of each studied population, we included interactive plots in Github (https://github.com/Schlebusch-lab/Sahel_study).

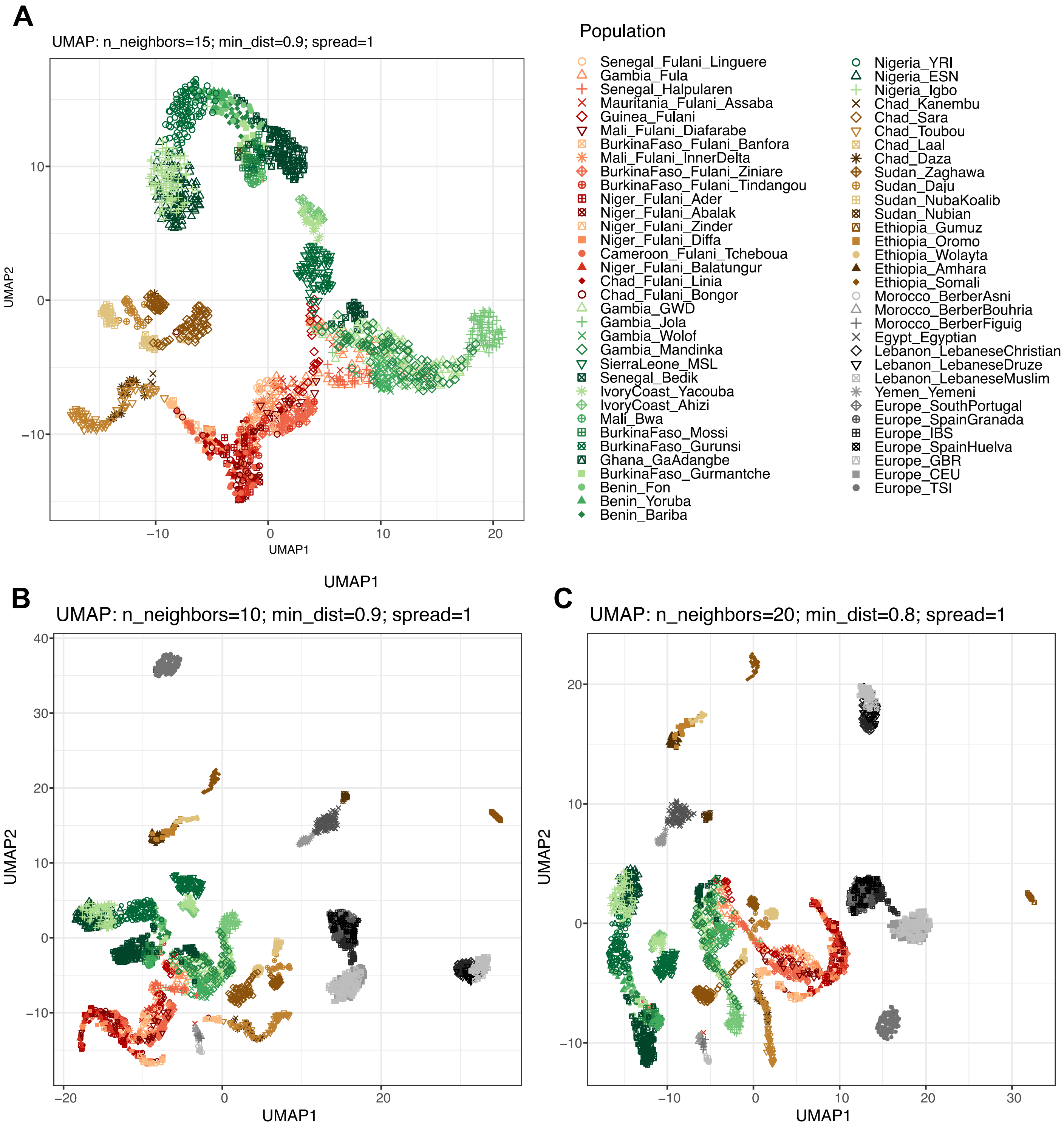


**Figure S6. Genome-wide diversity of the Fulani groups in the context of a broader genetic diversity of reference groups.** Figure showing PCA-UMAP combining the information of the first principal components of the PCA estimated for all the populations included in the Fulani-World dataset (**Table S2**). We performed PCA-UMAP for (A) western and central African populations (using different parameters than in **Figure 1F-1G**), and for (B, C) all comparative populations using different parameters. To avoid sample size bias (**Figure S4**), we first computed PCA for reference populations and a downsampled set of 36 Fulani individuals from all studied Fulani populations and we then projected onto the PCA the remaining Fulani samples. Geographical locations of the studied populations were included in **Figure S3A**.

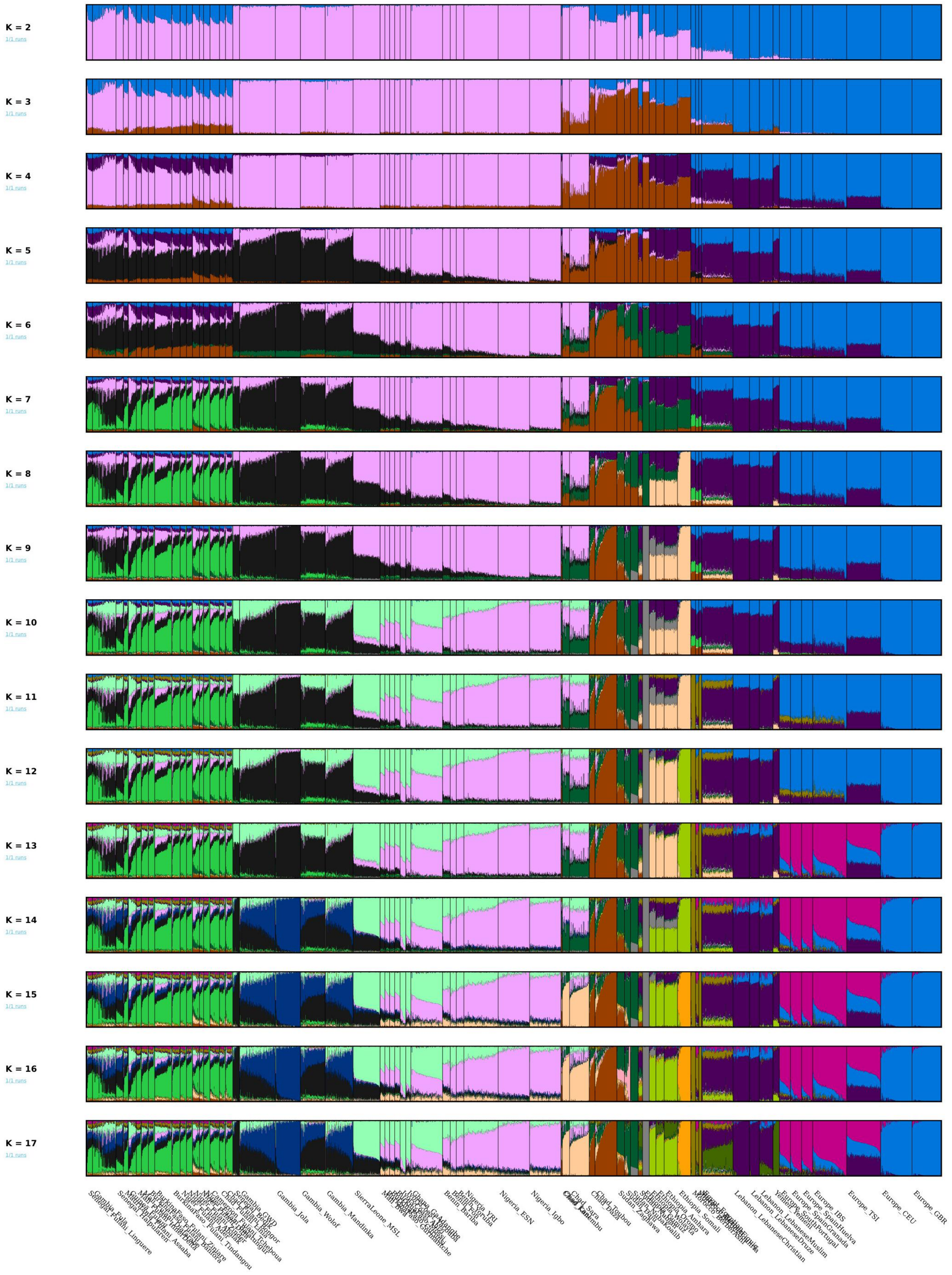


**Figure S7. ADMIXTURE analysis from K=2 to K=17.** ADMIXTURE results for K=7 showed lowest CV-error (**Figure S8D**), where clusters are assign to different putative ancestries: Fulani-related ancestry with the green component; Niger-Congo Atlantic with the black component; Niger-Congo Volta-Niger with the pink component; Nilo-Saharan Tubu with the brown component; Nilo-Saharan Gumuz with the dark green component; Afro-Asiatic with the purple component; and Indo-European with the blue component.

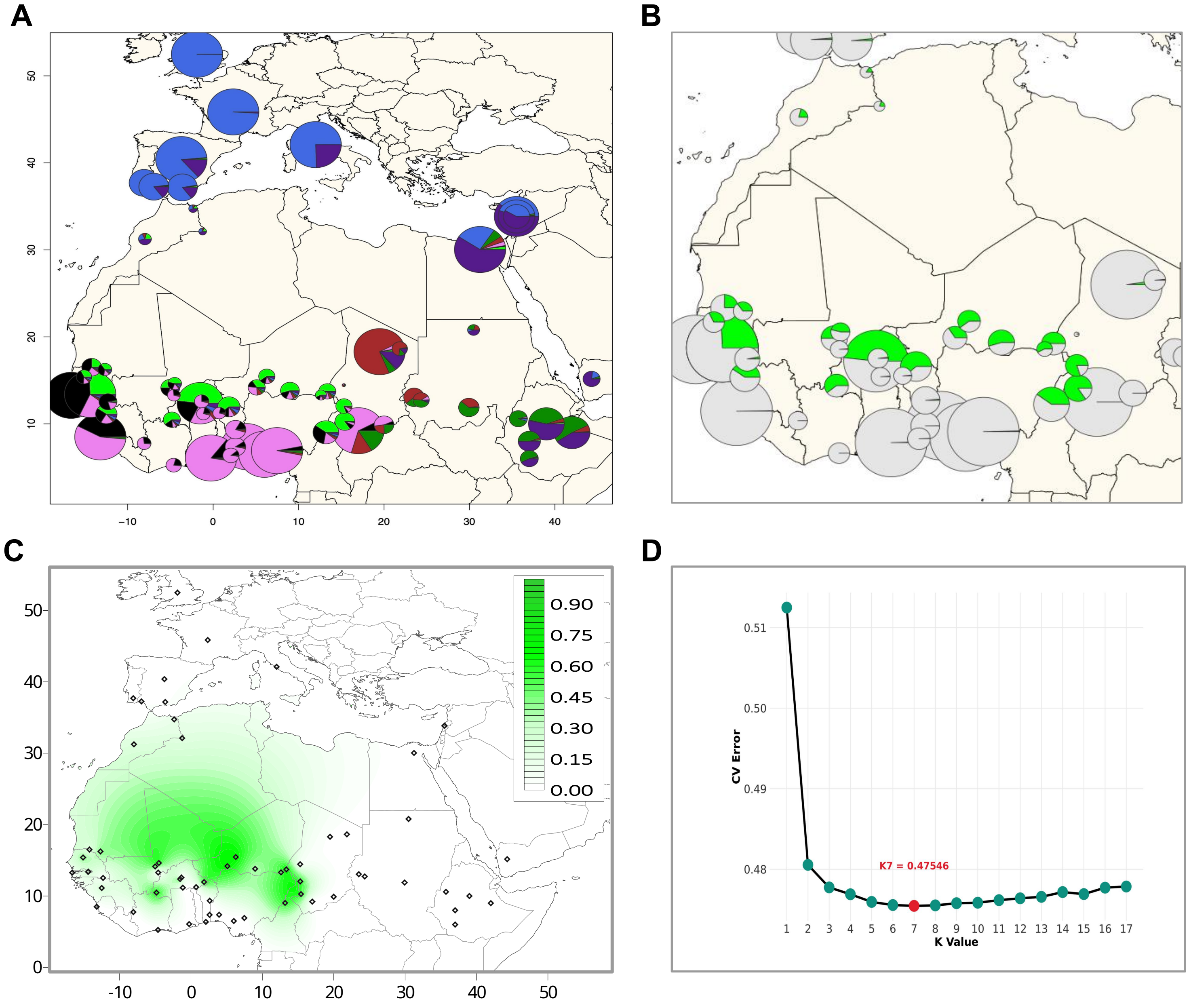


**Figure S8. ADMIXTURE results at K=7 using the projection mode.** (A) Figure showing pie charts to highlight the ADMIXTURE results at K=7 for all studied populations included in the Fulani-World dataset; and (B) to highlight the distribution of the Fulani-related component (in green) among African populations (other components are depicted in gray). (C) Contour map to visualize the distribution of the Fulani-related component across the African continent. (D) Average values of the cross-validation (CV) test for each K inferred in the ADMIXTURE analysis from K=2 to K=17 (**Figure S6**). Inferred components in the ADMIXTURE analysis at K=7 (A) could be assigned to different the following ancestries: Fulani-related ancestry with the green component; Niger-Congo Atlantic with the black component; Niger-Congo Volta-Niger with the pink component; Nilo-Saharan Tubu with the brown component; Nilo-Saharan Gumuz with the dark green component; Afro-Asiatic with the purple component; and Indo-European with the blue component. Estimated averages and standard deviations for each component were included in **Table S7**.

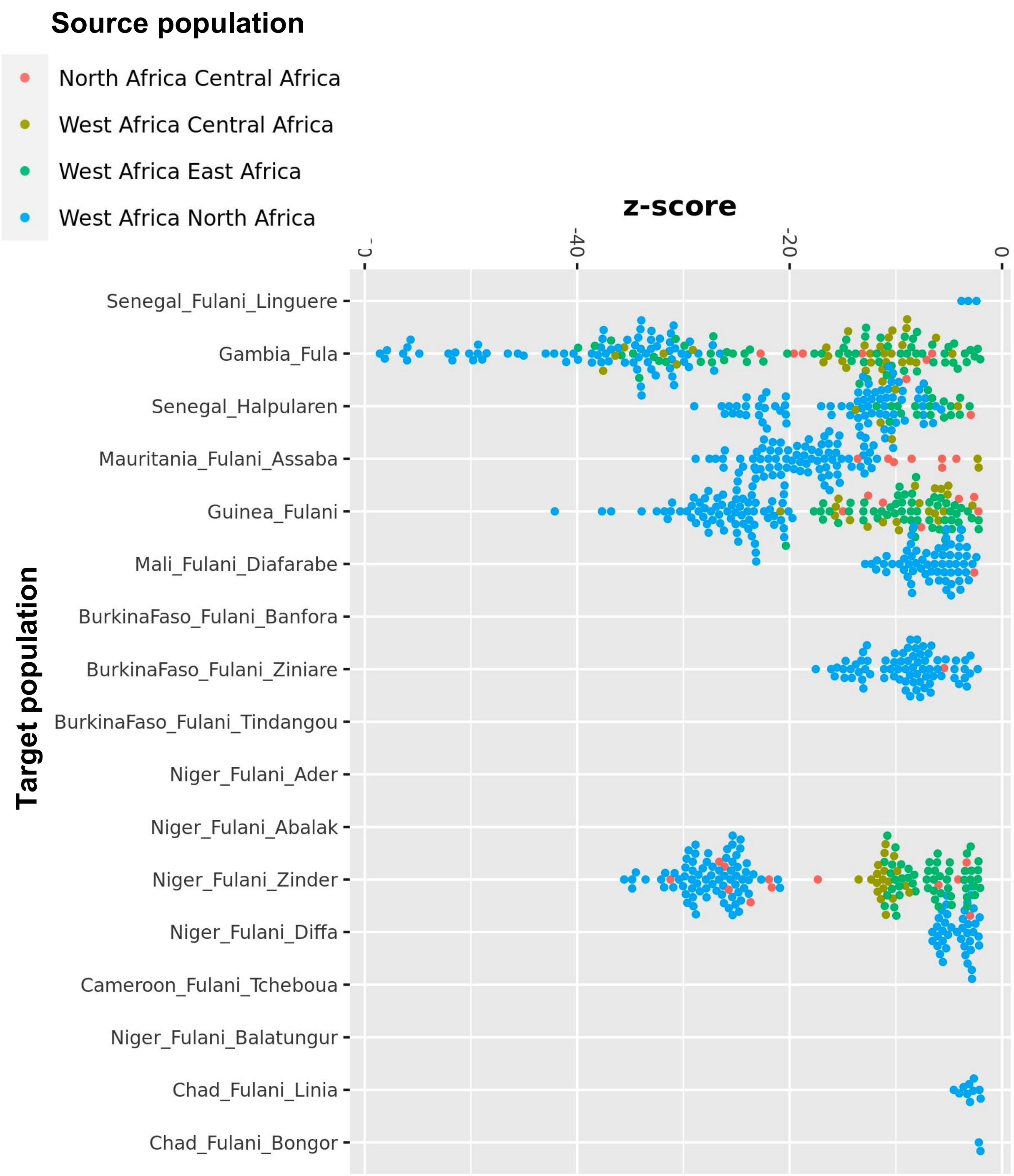


**Figure S9. Admixture tests using *f3*-statistics analyses.** Dots represent comparisons for pairs of source populations and specific Fulani populations (y-axis). The significance of *f3*-statistics is represented by Z-scores on x-axis. Only values below -2 were included in the figure. All estimated values were included in **Table S8**.

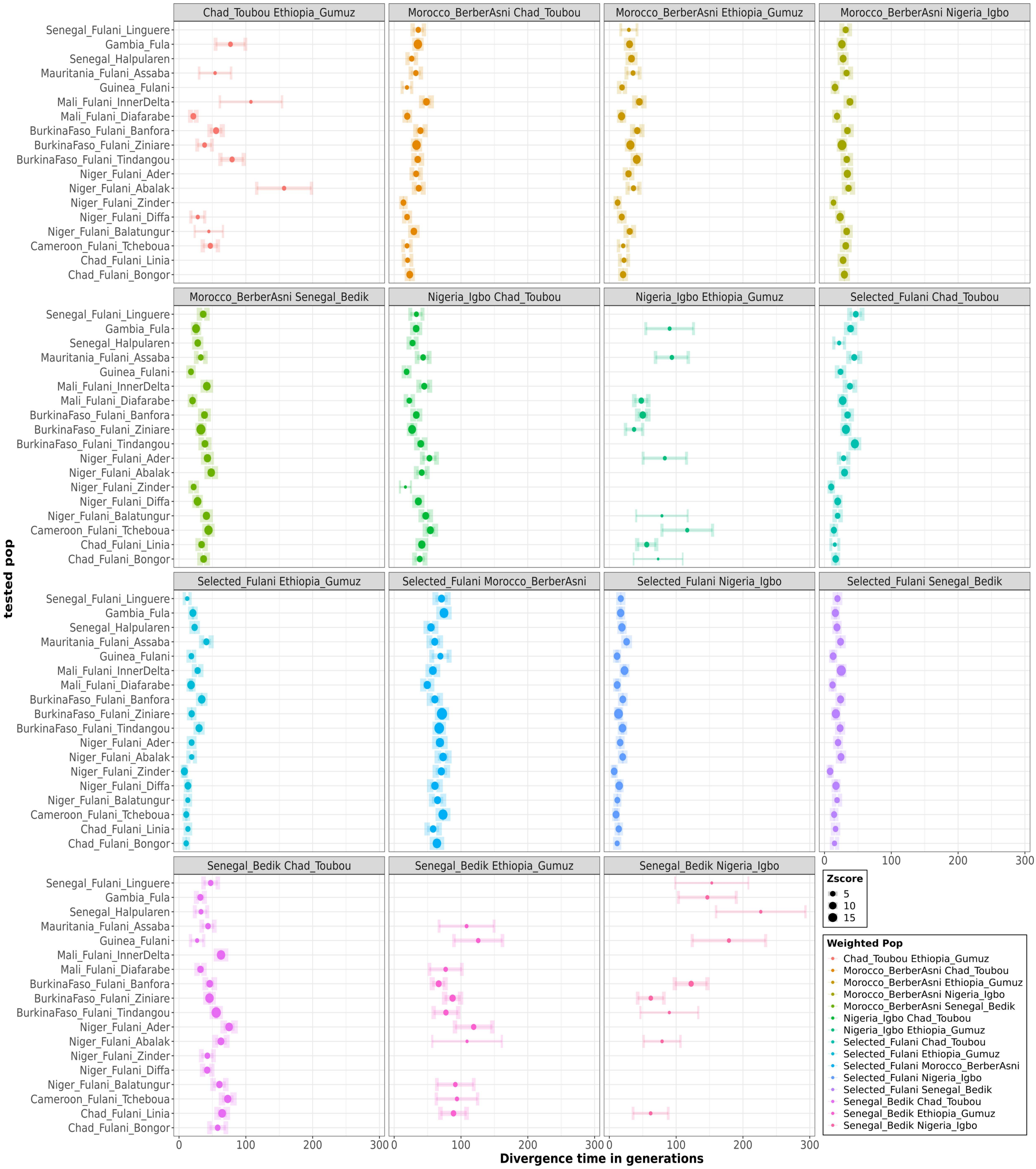


**Figure S10. Admixture timing resutls estimated using MALDER.** Admixture dates were estimated for multiple reference test divergence time among Fulani populations. Dots represent divergence times (in generations) delimited by line interval and colored by divergence source and sized by Z-score. B. Admixtures LD time grouped when Selected Fulani was inferred.

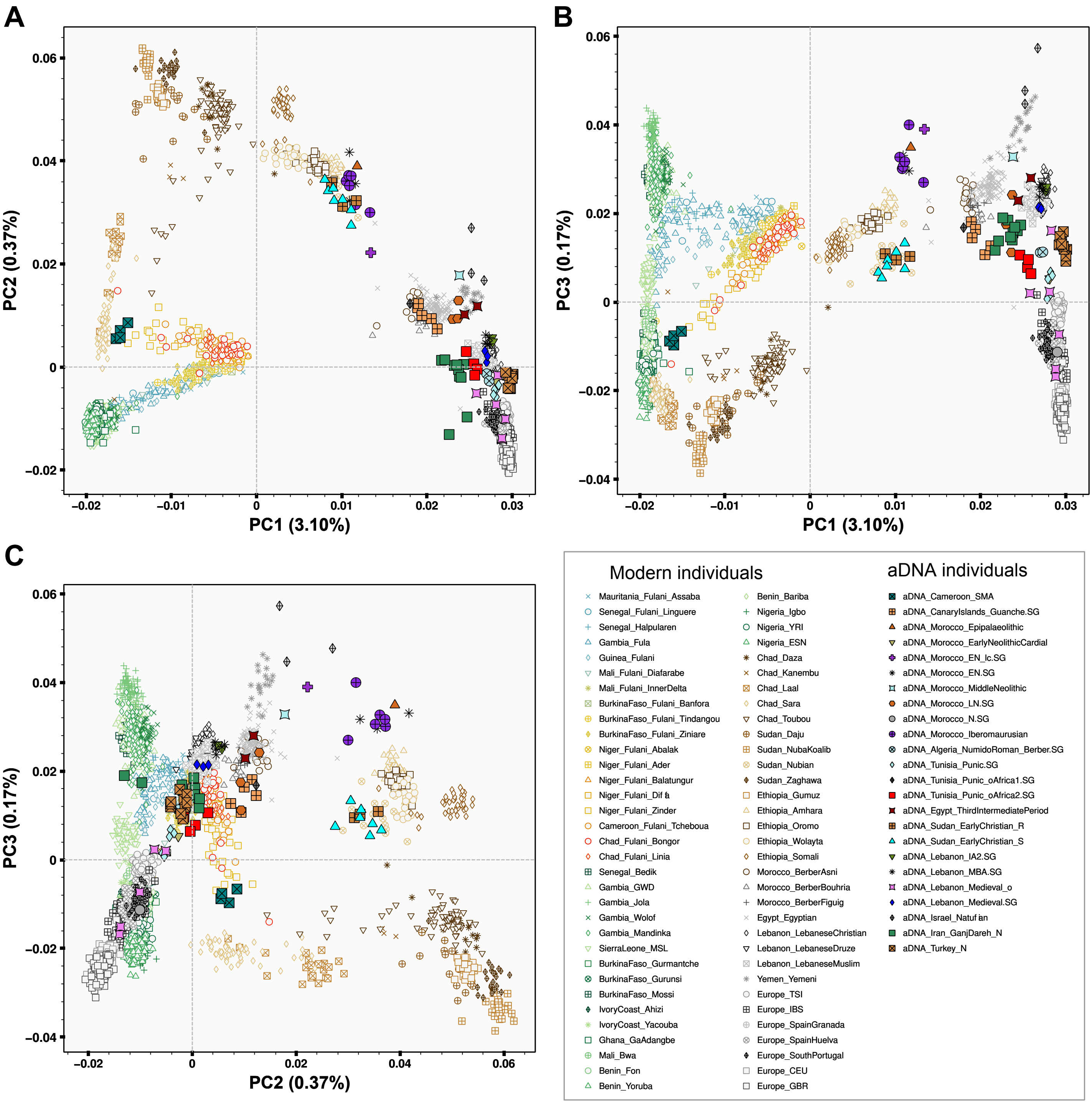


**Figure S11. PCA on the basis of modern and ancient individuals.**
Figure showing PC1 to PC3 obtained using smartPCA to project 91 ancient samples onto the background of present-day African populations (using "YES" option for the following parameters: allsnps, lsqproject, newshrink, and killr2). Markers of ancient samples were filled with different colors, while markers of modern populations have the same shape and border color than in **Figure S4**, but were filled with white. Details about the studied aDNA samples were included in **Table S3**. To better visualize the results of each studied population, we included interactive plots in Github (https://github.com/Schlebusch-lab/Sahel_study).

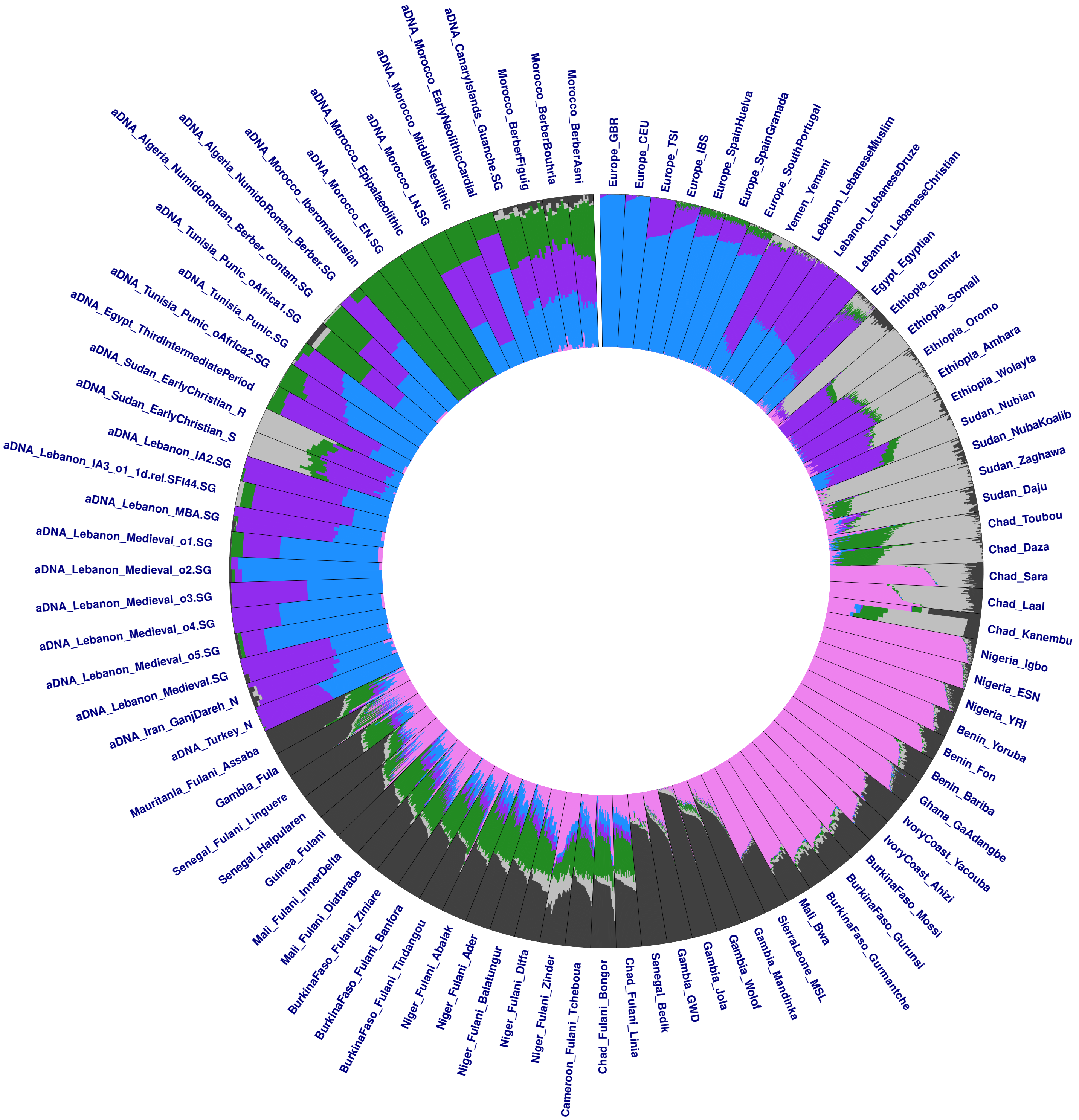


**Figure S12.** ADMIXTURE analyses results at K=6 on the basis modern and aDNA individuals, using the projection mode for selected Fulani individuals. For a better comparison, the width of each population was set to be equal regardless of its sample size.

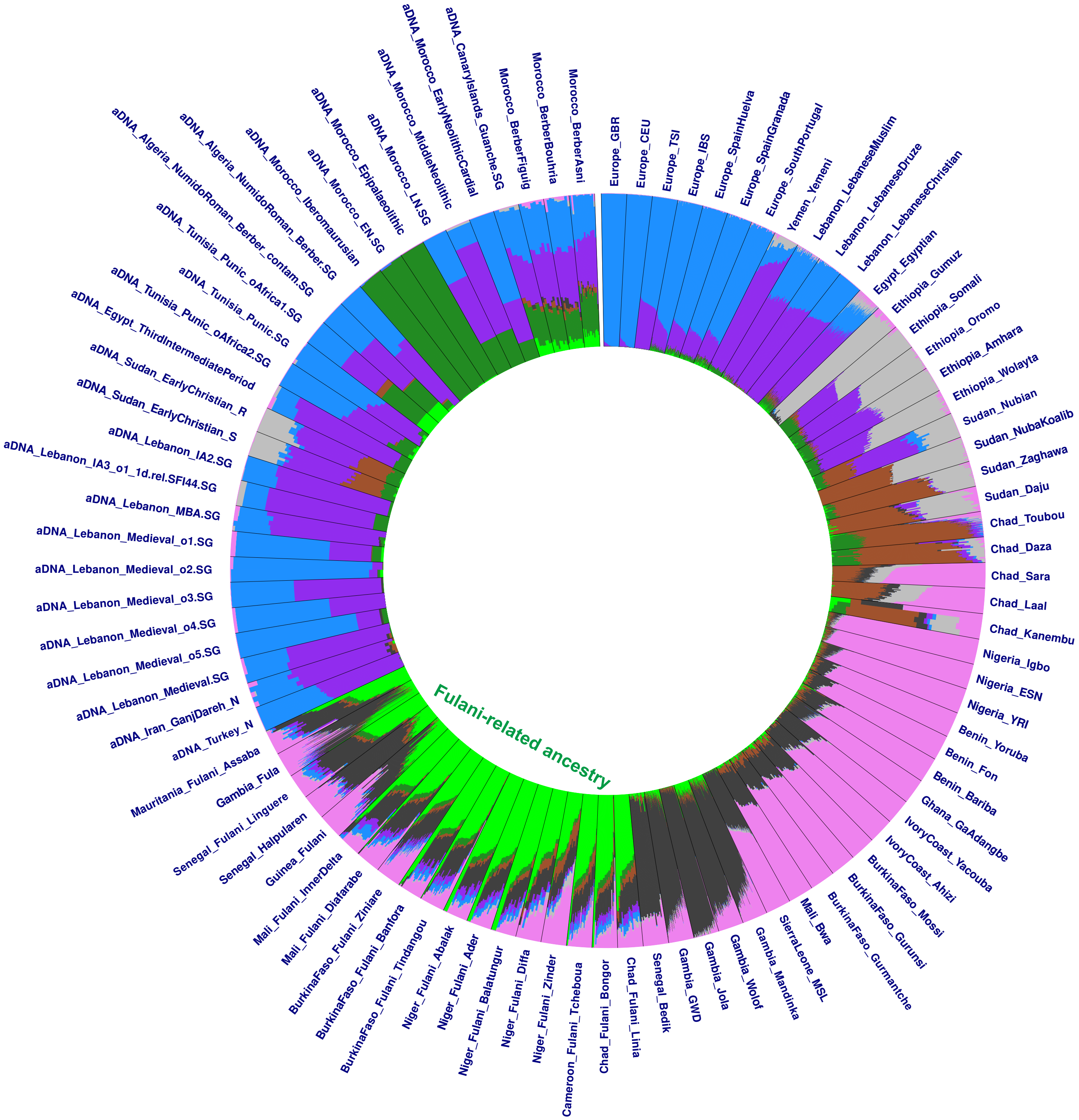


**Figure S13.** ADMIXTURE analyses results at K=8 on the basis modern and aDNA individuals, using the projection mode for selected Fulani individuals. For a better comparison, the width of each population was set to be equal regardless of its sample size.

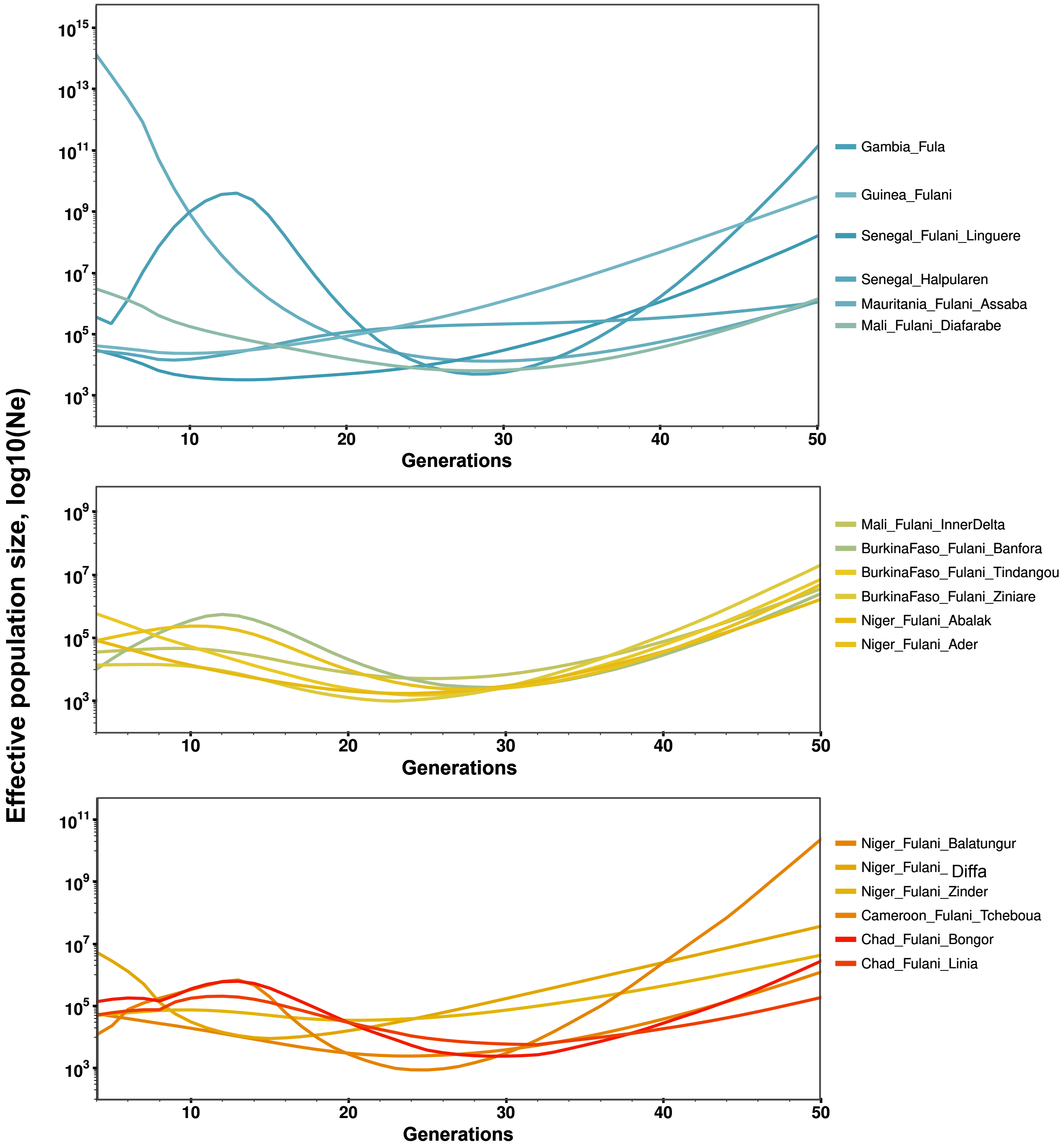


**Figure S14. Estimated effective population sizes for Fulani populations.** Effective population sizes ($N_\epsilon$) in studied Fulani populations for the last 50 generations estimated using IBDNe. Figure showing the results for Fulani populations from the western (top), west-central (middle), and central (bottom) region in the Sahel belt. These three groups were selected based on their west-east position and by including up to six populations for each group. All Fulani populations together were presented in **Figure 4A**. Estimated $N_\epsilon$ and two-tailed 95% confidence interval were included in **Table S11**. To better visualize the results of each studied population, we included interactive plots in Github with different options of zooming (https://github.com/Schlebusch-lab/Sahel_study).

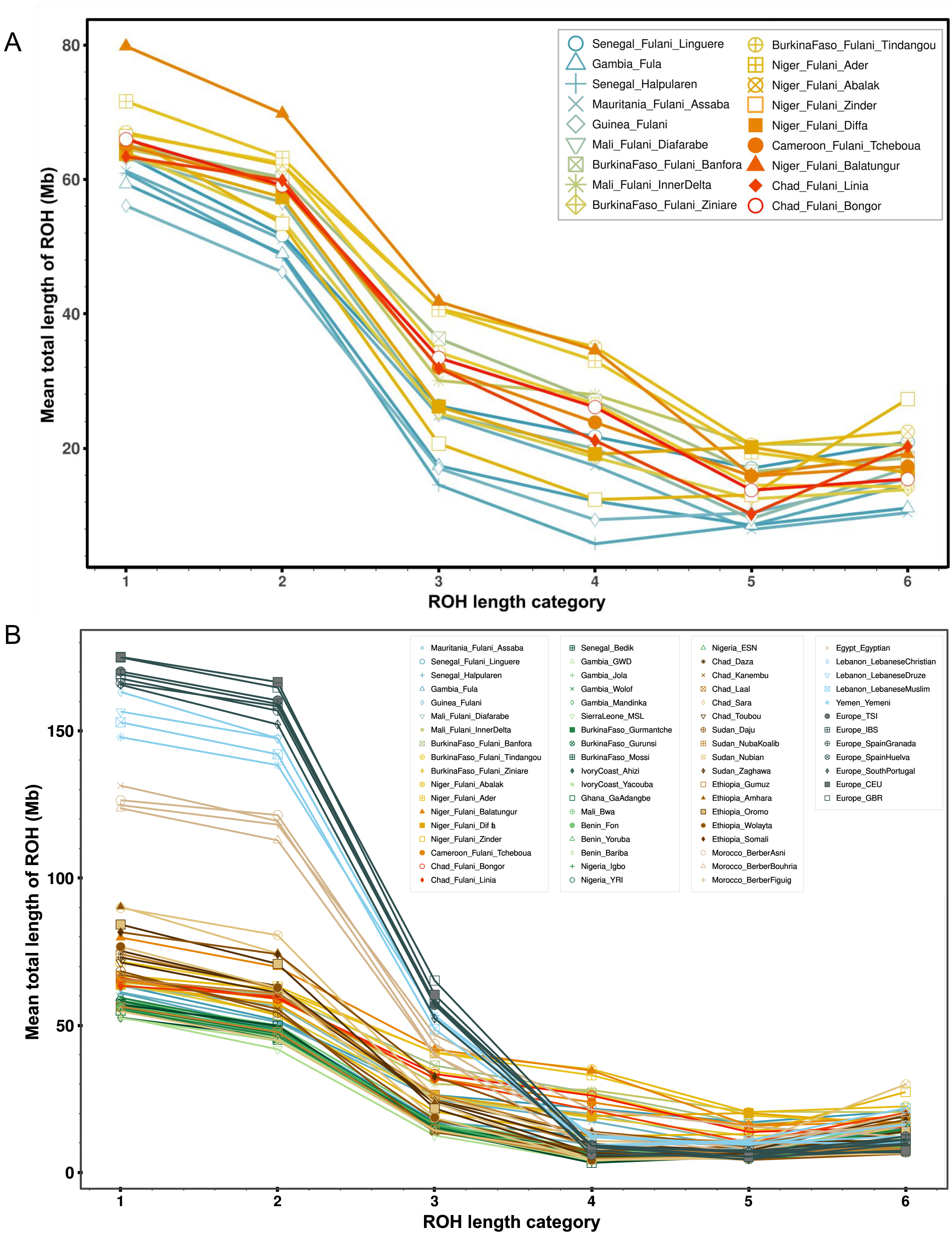


**Figure S15. Categories of ROH length on the basis of the studied populations.**
Figure showing averages in each studied population included in the (A) Fulani-Only dataset and (B) the Fulani-World dataset. For each category of ROH length includes the following lengths: class 1 for [0.3-0.5Mb); clas 2 for [0.5-1Mb); class 3 for [1-2Mb); class 4 for [2-4Mb); class 5 for [4-8Mb); and class 6 for [8-16Mb). To better visualize the results of each studied population, we included interactive plots in Github with different options of zooming (https://github.com/Schlebusch-lab/Sahel_study).

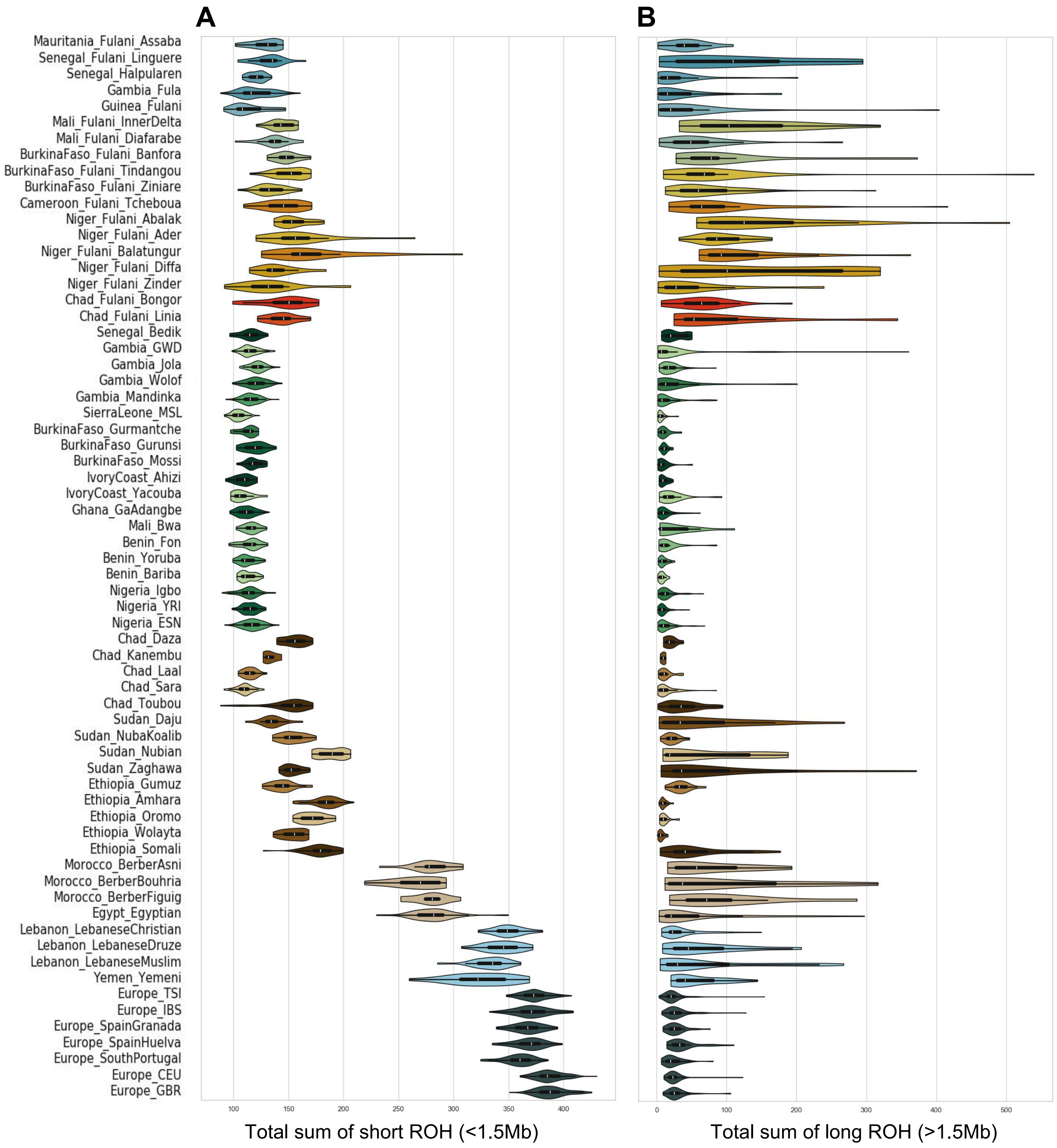


**Figure S16. ROH results for each studied population.** Figure showing violin plots for (A) the total sum of short ROH (<1.5Mb), and (B) the total sum of long ROH (>1.5Mb) estimated for each Fulani (top) and reference worldwide population. Mean and standard deviation values were included in **Table S12**.

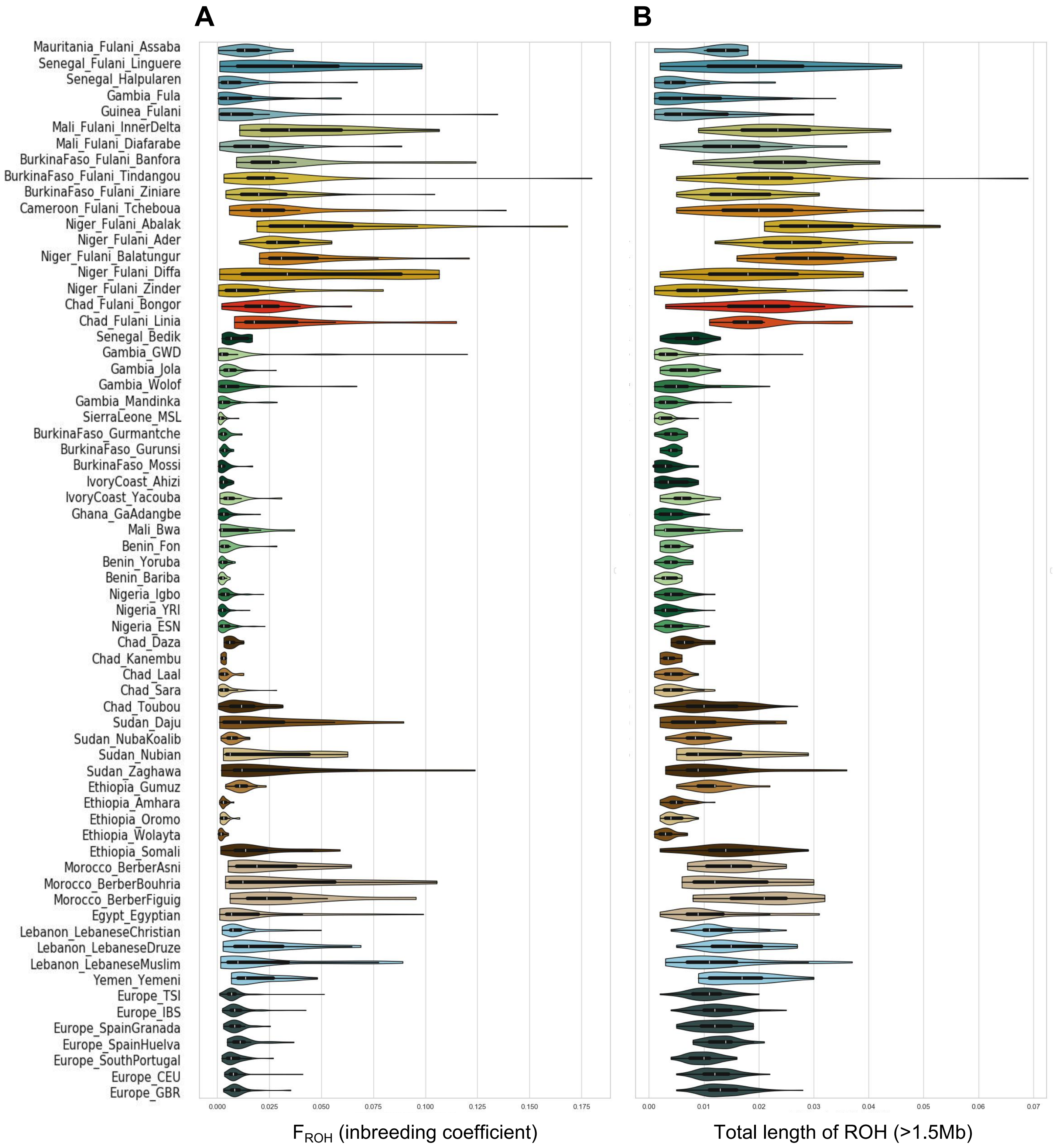


**Figure S17. ROH results for each studied population.** Figure showing violin plots for (A) genomic inbreeding coefficient based on ROH (or $F_{ROH}$) and (B) the total length of ROH longer than 1.5 Mb. $F_{ROH}$ measures the actual proportion of the autosomal genome that is autozygous, and was estimated based on the total sum of ROH>1.5 Mb divided by the total length of the autosomal genome (3 Gb). Mean and standard deviation values were included in **Table S12**.

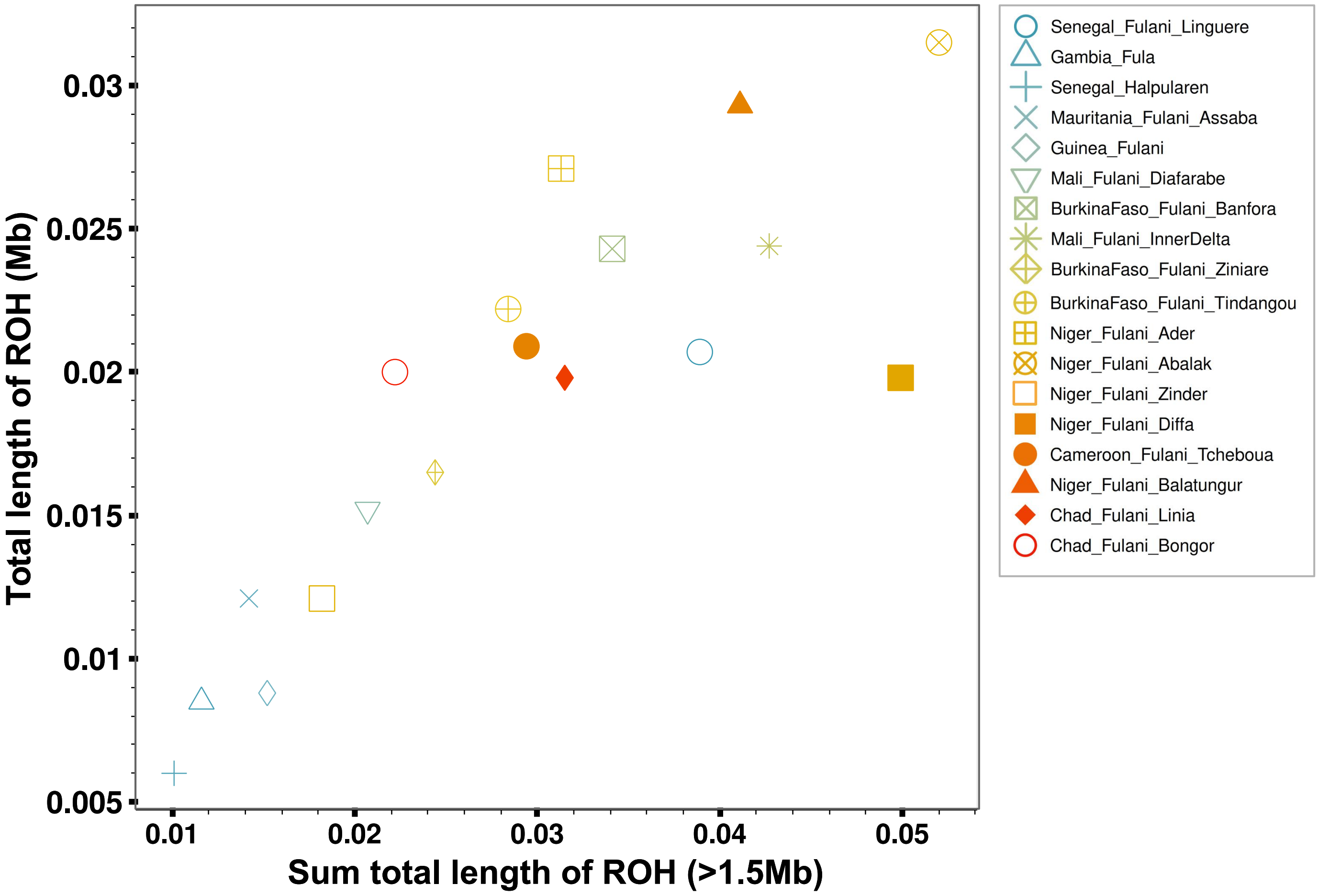


**Figure S18. Mean values of ROH for each Fulani population.** Figure comparing the mean values of the sum total length of ROH (>1.5Mb) and the total length of ROH (>1.5 Mb) for each studied Fulani population.

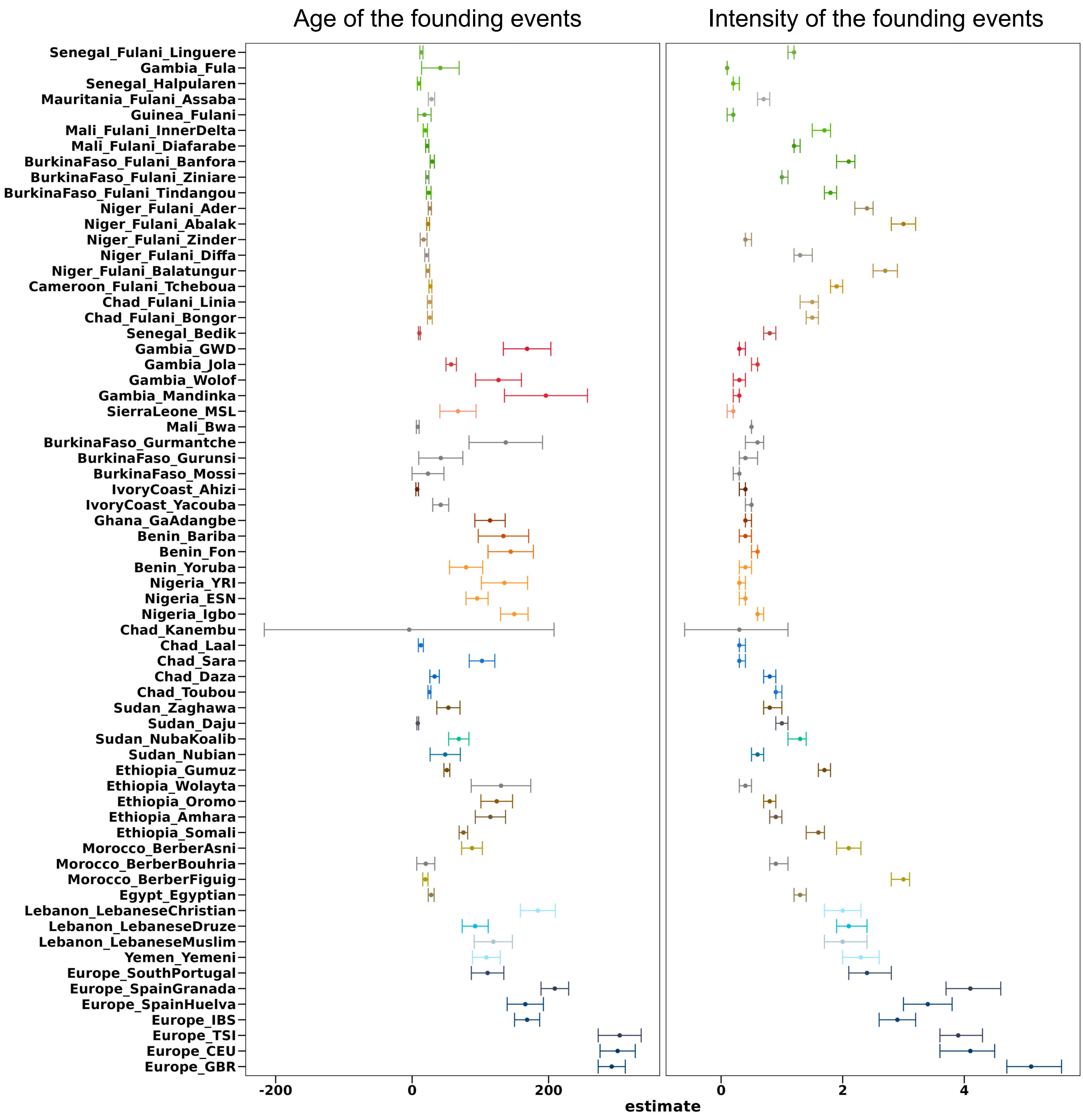


**Figure S19.** ASCEND results inferred for all the populations included in the Fulani-World dataset. Figure showing the mean values of the estimated founder ages in generations (left) and the estimated founder intensities (right) for each studied population and with their respective 95% confidence intervals. Estimated values were also included in **Table S12**.

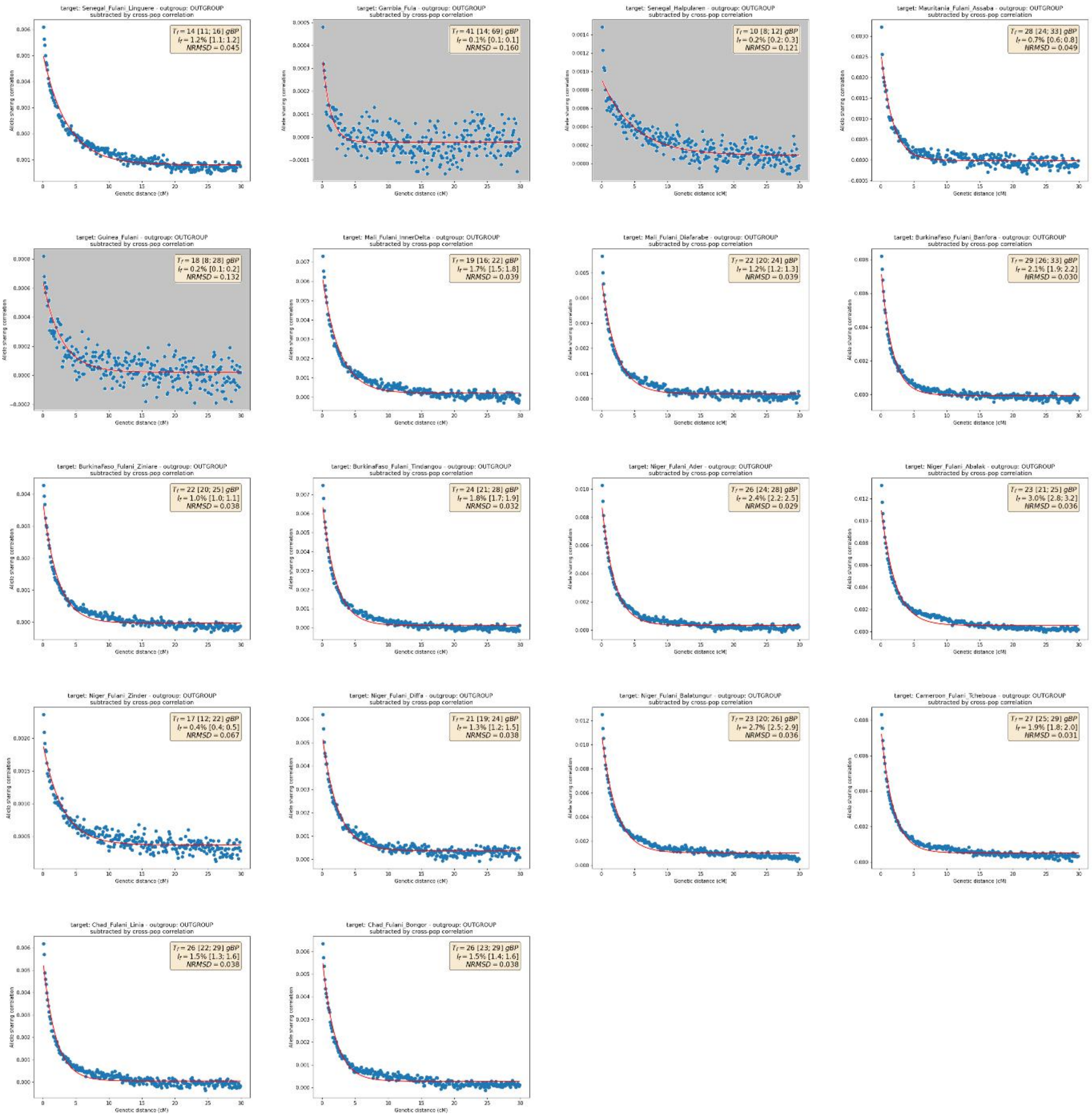


**Figure S20. ASCEND results for studied Fulani populations.** The plot of the allele sharing correlation decay curve (blue points) along with the fitted exponential model (red line). In the top-right corner: the estimates of founder age ($T_f$) and intensity ($I_f$) with their associated 95% confidence intervals within brackets as well as the NRMSD. We display plot only for studied Fulani populations. To assess the validity of the exponential fit, we estimated the normalized root-mean-square deviation (NRMSD) between the empirical allele-sharing correlation values and the fitted ones, and we plotted the correlations between empirical and theoretical decay curves.
